# CDK4/6 Inhibition Reverses MEIS2 Suppression of CRL4^CRBN^ to Enhance Immunomodulatory Drug Therapy in Multiple Myeloma

**DOI:** 10.1101/2025.10.14.681840

**Authors:** Xiangao Huang, David Jayabalan, Maurizio Di Liberto, Zhengming Chen, Scott A. Ely, Tomer M. Mark, Gang Lin, Ruben Niesvizky, Selina Chen-Kiang

## Abstract

MEIS2 was identified biochemically as a substrate of cereblon (CRBN), a receptor of the CRL4^CRBN^ E3 ubiquitin ligase required for the anti-myeloma activity of immunomodulatory drugs (IMiD)s and CELMoDs. However, its function in myeloma is unknown. We discovered that MEIS2 is aberrantly expressed in bone marrow myeloma cells (BMMC)s, and that high *MEIS2, CDK4 or CDK6* and low *CRBN* expression predisposes patients to inferior overall survival in IMiD therapy. Inhibition of CDK4/6 (CDK4/6i) reprograms BMMCs for IMiD vulnerability *ex vivo* that mimics patient’s response to IMiDs. Mechanistically, CDK4/6i both rapidly accelerates the displacement of MEIS2 from CRBN by IMiD and destabilizes the MEIS2 protein while increasing the CRBN protein in cooperation with IMiD and CELMoD. This enhances CRL4^CRBN^ ubiquitination of IKZF3 and IKZF1 for degradation that exacerbates the loss of IRF4 to relieve IRF7 for induction of the interferon response, culminating in TRAIL-mediated apoptosis. Additionally, MEIS2 promotes BCMA expression and antagonizes repression of BCMA by IMiD and CELMoD for survival of myeloma cells. Thus, CDK4/6i reverses MEIS2 inhibition of CRL4^CRBN^ in cooperation with IMiD and CELMoD, and mitigates MEIS2-mediated BCMA signaling for survival, suggesting targeting MEIS2 and CDK4/6 as a new strategy to advance immunomodulatory drug therapy in multiple myeloma.

## Introduction

The immunomodulatory drugs (IMiDs) lenalidomide (Len) and pomalidomide (Pom) are standard of care for multiple myeloma (MM), the neoplasm of plasma cells. Despite a high overall response rate, patients treated with these agents alone rarely achieve complete remission; however, when given with proteasome inhibitors or antibodies to CS-1 or CD38, some MM patients enjoy robust, deep, and long complete responses ^1, 2, 3, 4, 5^. Following the discovery that thalidomide bound to cereblon (CRBN) ^6^, a substrate receptor of the CUL4-RBX1-DDB1-CRBN (CRL4^CRBN^) E3 ubiquitin ligase, the mechanism of IMiD action came to light as Len and Pom also bound to CRBN directly ^7^. CRBN is required for IMiD’s anti-tumor activity ^8, 9^. Binding of IMiD to CRBN promotes recruitment of IKZF1 and IKZF3 to CRL4^CRBN^ for ubiquitination-proteasome degradation in myeloma cell lines and primary myeloma cells ^10, 11^. This leads to downregulation of a direct target IRF4 ^10^ required for survival of myeloma cell lines^12^. Owing to its high affinity for CRBN ^13^, the second generation IMiD **c**ereblon **E**3 **l**igase **mo**dulatory **d**rug (CELMoD) is more potent than Len or Pom in myeloma therapy, even in Len-refractory patients ^13, 14^. These findings reinforce the critical importance of modulating CRBN for clinical response to IMiDs,

Little is known, however, about regulation of CRBN. MEIS2, a transcription factor of the three amino-acid loop extension family ^15^ involved in various aspects of vertebrate development ^16, 17, 18, 19^ and oncogenesis of acute myeloid leukemia ^20^, was identified as an endogenous substrate of CRBN by human protein microarray ^21^. MEIS2 competes with Len for binding to the same domain of CRBN *in vitro* and in various cell lines ^21^, implicating a role for MEIS2 in regulating the function of CRL4^CRBN^. This remains to be proven, and the role of MEIS2 in IMiD therapy is unknown.

Disease progression and therapy resistance in MM are frequently associated with proliferation of primary bone marrow myeloma cells (BMMCs) stemming from coordinated overexpression of cyclin D1 and CDK4, or cyclin D2 and CDK6 (CDK4), that drives cell cycle progression through early G1 ^22^. Selective inhibition of CDK4/CDK6 by palbociclib (PD 0332991) ^23^ arrested BMMCs in early G1 *ex vivo* ^24^ and suppressed myeloma tumor growth in xenografts and the immunocompetent 5T mouse model of myeloma ^24, 25, 26^. Moreover, induction of prolonged early G1 arrest (pG1) by sustained CDK4/6 inhibition (CDK4/6i) reprogrammed myeloma cells for killing by partner agents such as dexamethasone and bortezomib *ex vivo* and *in vivo* ^24, 25, 26^. In line with these earlier findings, CDK6 was identified as a marker for IMiD resistance by proteasome profiling, and inhibiting CDK4/6 or degrading CDK6 by PROTAC augmented IMiD’s anti-MM activity in MM cell lines and xenografts ^27, 28^. However, the precise mechanism by which CDK4/6i sensitizes MM cells to IMiDs and the role of MEIS2 and CRBN await investigation. Whether CDK4/6i augments CELMoD’s anti-myeloma activity is also unknown.

To address these questions, we have now discovered that expression of high *MEIS2, CDK4 and CDK6* and low *CRBN* in BMMCs predispose myeloma patients to inferior overall survival in IMiD therapy. CDK4/6i primes BMMCs for IMiD vulnerability *ex vivo* in the context of the clinical response to IMiD. MEIS2 inhibits CRL4^CRBN^ and promotes survival by sustaining BCMA expression. CDK4/6i priming restores the CRL4^CRBN^ function by both disrupting MEIS2-CRBN association and reducing the MEIS2 protein while elevating the CRBN protein in cooperation with IMiD and CELMoD. This promotes apoptosis through the IKZF1/3-IRF4-IRF7-IFN-TRAIL pathway in conjunction with repressing BCMA by IMiD and CELMoD, suggesting inhibition of MEIS2 and CDK4/6 as a novel strategy to enhance IMiD and CELMoD therapy in multiple myeloma.

## Results

### Association of high MEIS2 expression with inferior overall survival in IMiD therapy

To investigate the role of MEIS2 in clinical response to IMiDs, we discovered that high *MEIS2* mRNA expression in bone marrow myeloma cells (BMMC)s before therapy strongly correlated with inferior overall survival (OS) (p =0.0383) in patients treated with IMiD monotherapy (n=37) in the Multiple Myeloma Research Foundation (MMRF) CoMMpass Study (n=859) (NCT01454297). Conversely, high *CRBN* expression positively correlated with improved OS (p = 0.0131) (Fig. 1a), as reported ^37^. Confirming the IMiD specificity, there was no association of *MEIS2* or *CRBN* expression with OS in patients treated with bortezomib-based therapy (n=112) (Fig. 1a). However, expression of low *CRBN* and high *CDK4*, *CDK6, E2F1 and CDK2* that drive cell cycle progression through G1 also correlated with inferior OS in the Lenalidomid-Bortezomib-Dexamethasone (Len-Bor-Dex) combination therapy (n=382) (Fig. 1a, b).

**Fig. 1.**
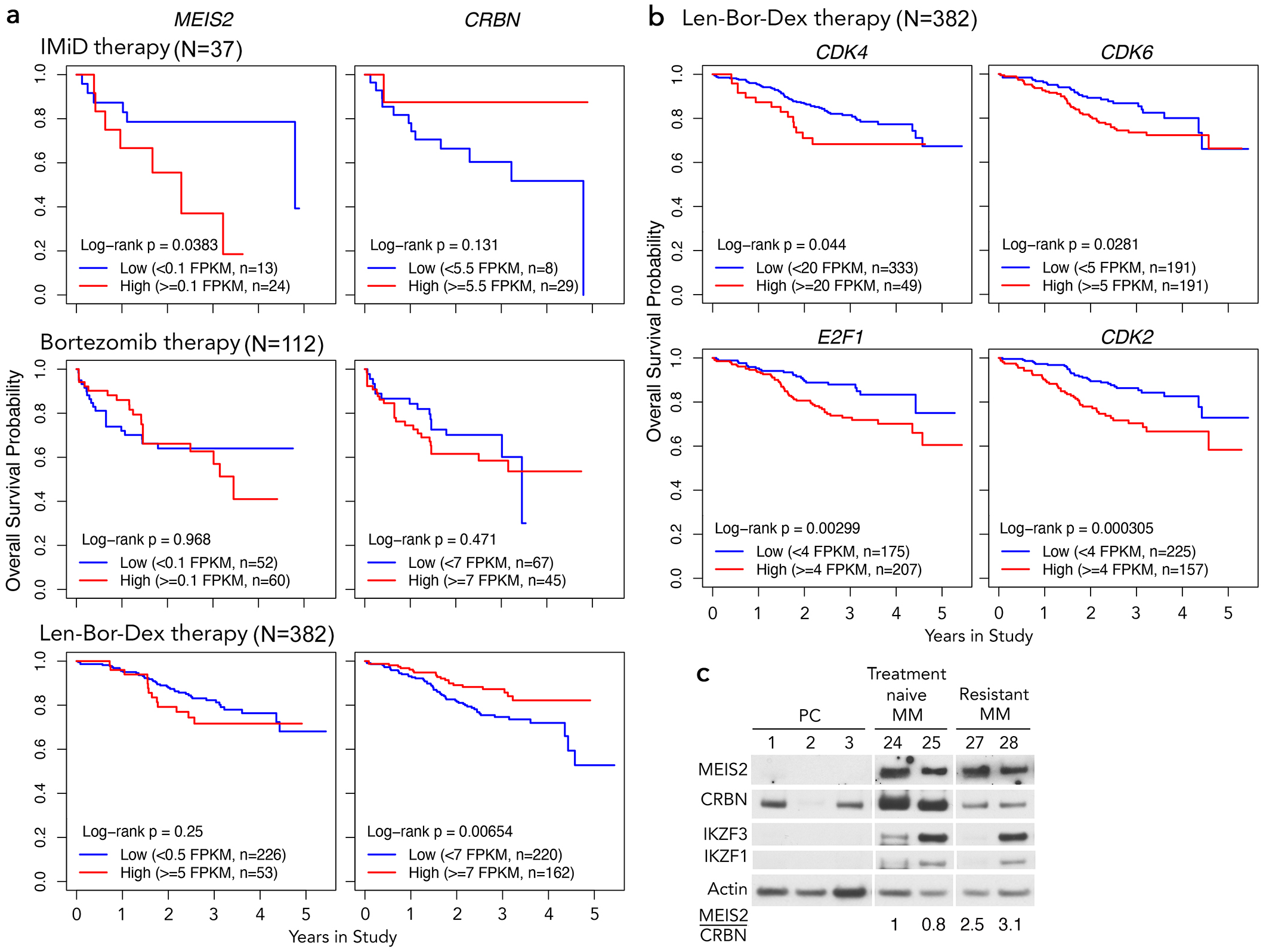
Association of high MEIS2 expression with inferior survival in IMiD therapy. **a** Kaplan-Meier plots of overall survival stratified by high and low expressions of *MEIS2* of *CRBN* for patients who *received only lenalidomide, pomalidomide or thalidomide with or without dexamethasone (IMiD therapy, N=37), bortezomib together with dexamethasone (Bortezomib therapy, N=112), or* lenalidomide-bortezomib-dexamethasone (Len-Bor-Dex therapy, N=382) from the MMRF CoMMpass study. **b** Kaplan-Meier plots of overall survival stratified by high and low expressions of the indicated cell cycle genes for patients who *received* Len-Bor-Dex therapy (N=382) from the MMRF CoMMpass study. **c** Immunoblotting of MEIS2, CRBN, IKZF3 and IKZF1 in normal plasma cells (PCs), bone marrow myeloma cells (BMMCs) from treatment naive or treatment refractory MM patients. Ratios of MEIS2 to CRBN are indicated.

*MEIS2* mRNA was highly expressed in BMMCs of some MM patients but barely detectable in normal plasma cells (PCs), in contrast to comparable mRNA expression of *CRBN, CUL4A, RBX1, DDB1* and genes of the IKZF family (Supplementary Fig. 1). Accordingly, the MEIS2 protein was expressed in BMMCs but not PCs (Fig. 1c). While the CRBN protein level varied in PCs, it was significantly higher in myeloma cells of treatment-naïve than resistant patients, which resulted in a 3-fold higher MEIS2 to CRBN ratio in resistance (Fig. 1c). As MEIS2 is an endogenous substrate of CRBN in myeloma cells ^21^, these results implicate a role for MEIS2 in inhibiting CRL4^CRBN^ to promote clinical resistance to IMiDs in concert with CDK4 or CDK6 dysregulation.

### CDK4/6i enhances IMiD killing of primary bone marrow myeloma cells

To address this possibility, we investigated whether CDK4/6i sensitizes freshly isolated BMMCs to IMiD in co-culture with HS-5 stromal cells ^38^ and cytokines, which sustains the cell cycle in BMMCs for the initial 10∼12 hours after isolation ^24^ (Fig. 2a). Inhibition of CDK4/6 (CDK4/6i priming) with palbociclib (Pal) for 6-8 h induced early G1 arrest in BMMCs, given the loss of CDK4/6-specific phosphorylation of Rb (pRb) (MM32) (Fig. 2b) and the downregulation of *E2F1* and its targets *CDC6, PCNA*, *TK1,* and *AURKB* ^39, 40^ (MM33, MM34)(Fig. 2c). Killing of BMMCs by Len was time- and dose-dependent, greater with CDK4/6i priming (Fig. 2d, e). The majority of BMMCs (14/19) were sensitive to Len, whether they were isolated from newly diagnosed or previously treated MM patients, such as MM2, MM5 and MM9 with 9, 5, and 5 prior therapies, respectively (Fig. 2f and supplementary Table 1). CDK4/6i priming enhanced Len killing in a patient-dependent manner based on two-way ANOVA, apart from MM12 that was exceptionally sensitive to Len (Supplementary Table 2). While similarly enhancing Pom killing (3/4), CDK4/6i priming did not confer susceptibility to BMMCs that were marginally sensitive, or refractory to Len (5/19) or Pom (MM15) (Fig. 2g, h).

**Fig. 2.**
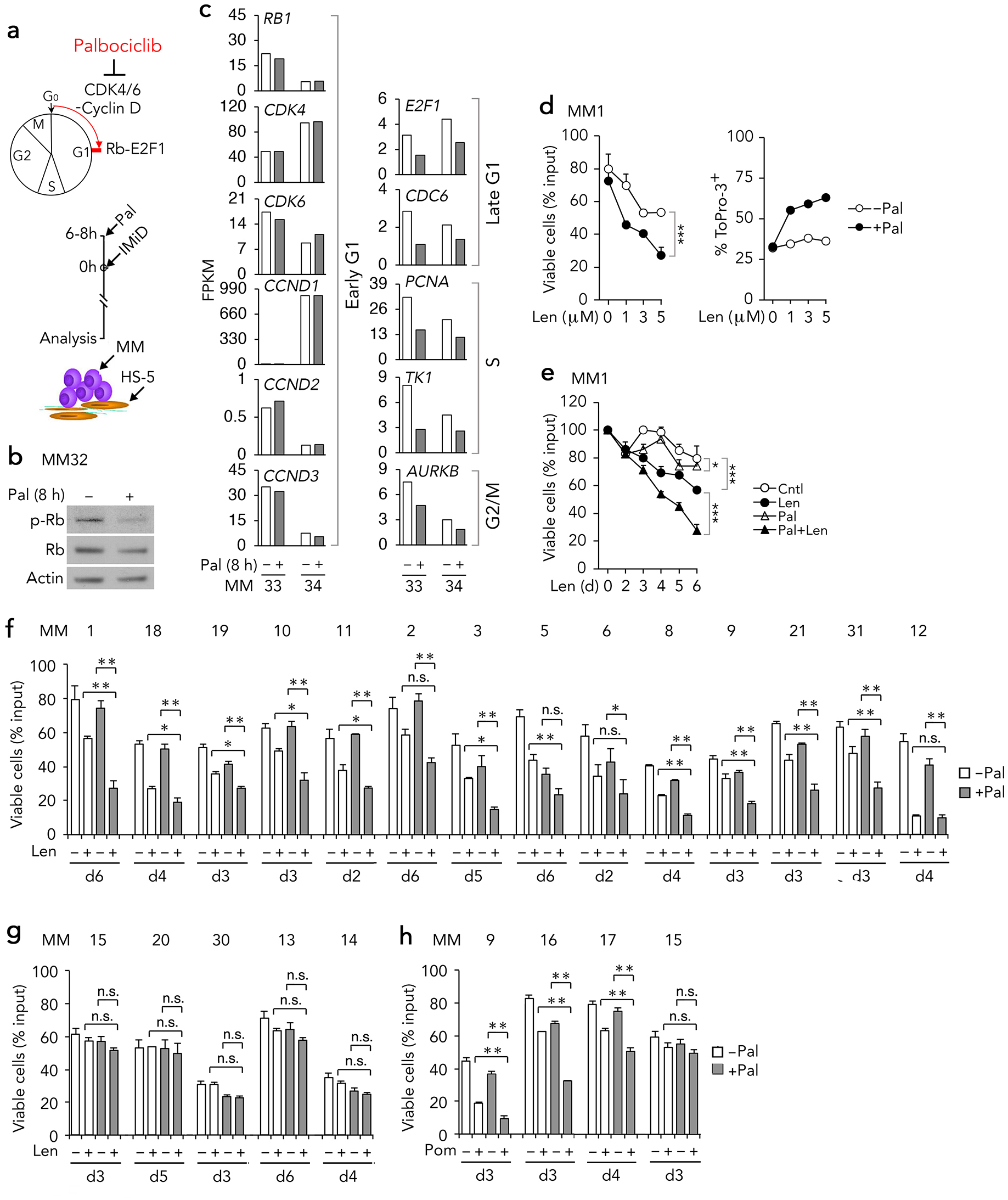
CDK4/6i enhances IMiD killing of primary myeloma cells. **a** Schema for induction of early G1 arrest by CDK4/6 inhibition with palbociclib (Pal) (upper) and an outline for sequential Pal and Len addition to co-culture of CD138^+^ BMMCs freshly isolated from patients (MM) with HS-5 bone marrow stromal cells (bottom). **b** Immunoblotting of Rb phosphorylated on serine 807-811 (p-Rb) in BMMCs with or without Pal (300 nM) for 8 h ex vivo. **c** RNA-Seq analysis of indicated genes in BMMCs treated as in (**b**). FKPM (fragments per kilobase of transcript per million mapped fragment) indicate RNA expression levels between different samples or genes. **d-e** Cell viability and cell death (To-Pro3+) in MM1 cultured with Len for 5 days at concentrations indicated (**d**) or at 3 µM for days indicated (**e**) with or without pal pretreatment for 6 h. **f-h** Cell viability in BMMCs cultured as in (**a**) for indicated days of Len (**f,g**) or Pom (3 μM) (**h**) treatment. The synergistic or additive effect of two-drug combination was determined by two-way ANOVA analysis of the square root transformed numbers of viable cells. The interaction and the pairwise comparison p-values were presented in the supplemental Table 1. *, p < 0.05; **, p < 0.01; n.s., not significant.

### Association of tumor intrinsic sensitivity with clinical response to IMiDs in myeloma

IMiD targets multiple cell types *in vivo*. Importantly, killing of BMMCs by IMiDs *ex vivo* (Fig. 2f, h) correlated with the clinical response to the first post-biopsy Len- or Pom-based therapy (referred to as IMiD therapy) in individual patients (n=16), ranging from complete response (CR) to very good partial response (VGPR), partial response (PR) and stable disease (SD) (Supplementary Table 1). Conversely, refractory to IMiDs in BMMCs *ex vivo* (n=5) (Fig. 2g, h) recapitulated resistance to IMiD therapy before biopsy in the 4 evaluable patients (MM 13, 15, 20, 30) (Supplementary Table 1). Thus, IMiD killing of primary myeloma cells in our reconstituted bone marrow stromal culture recapitulates the immediate clinical response to IMiD therapy in individual patients, indicating that the tumor intrinsic sensitivity is a major determinant for the clinical response to IMiDs in MM, and may be enhanced by CDK4/6 inhibition.

### CDK4/6i reduces MEIS2 and increases CRBN to restore CRL4^CRBN^ activity for IMiD killing

To investigate the underlying mechanism in primary myeloma cells, we discovered that CDK4/6i and Len each reduced the MEIS2 protein, and CDK4/6i also increased the CRBN protein (2-fold) greater when combined with Len (4-fold) by 48h in MM10 (Fig. 3a). Consequently, the MEIS2 to CRBN ratio was lowered by Len and CDK4/6i, alone and cooperatively, concurrent with depletion of IKZF1, IKZF3 and IRF4 proteins (Fig. 3a). MEIS2 or CRBN mRNAs did not change, suggesting opposing posttranscriptional regulation (PTR) of MEIS and CRBN proteins by CDK4/6i priming and Len. *IKZF1* and *IKZF3* mRNA also did not vary, but *IRF4* mRNA was downregulated by Len as expected of a IKZF1/3 target, and not by CDK4/6i, indicating it reduced the IRF4 protein by posttranscriptional mechanisms

**Fig. 3.**
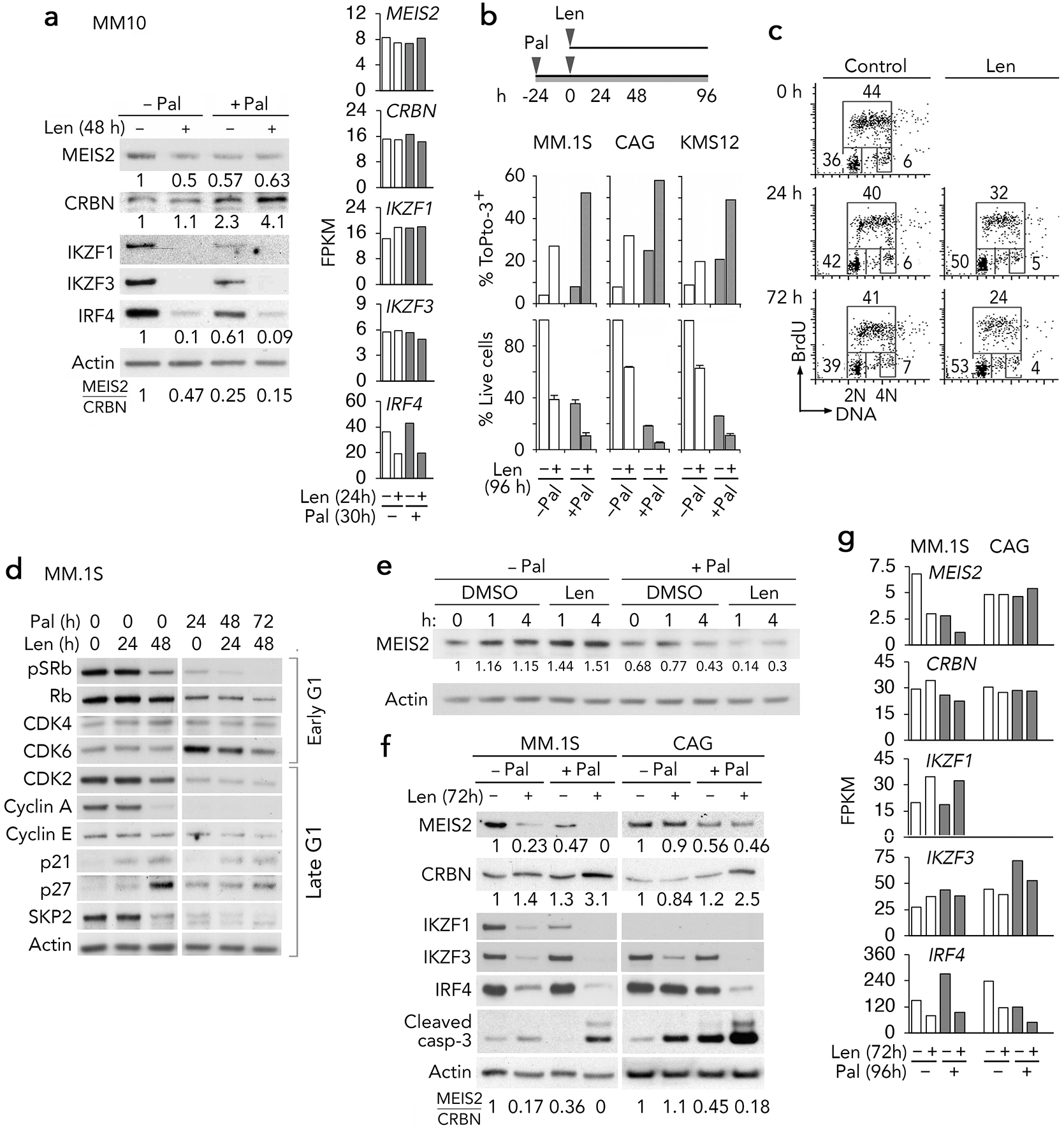
CDK4/6i reduces MEIS2 and increases CRBN to restore CRL4^CRBN^ activity for IMiD killing. **a** Immunoblotting of indicated proteins (left) and RNA-seq analysis of indicated genes (right) in BMMCs (MM10) cultured with Len for 48 h with or without Pal pretreatment for 6 h. Levels of MEIS2 and CRBN proteins were normalized to Actin and calculated against the untreated control (lane 1). The ratios of MEIS2 to CRBN protein were shown at the bottom (right). **b** Cell death determined by To-Pro3 staining and live cells determined by trypan blue exclusion staining after 96 h of Len treatment with or without 24 h Pal pretreatment. **c** FACS analysis of BrdU uptake in 30 minutes and DNA content in MM1.S cells cultured with Len for hour (h) indicated. The numbers indicate the percentage of cells in G1(2N), S (<2N<4N) or G2/M (4N) phase of the cell cycle. **d** Immunoblotting of indicated proteins in MM.1S cells treated with Pal (300 nM) and Len (3 µM daily) for times indicated. **e** Immunoblotting of MEIS2 protein in MM.1S cells treated with Len (30 µM) or DMSO for 1 or 4 h with or without Pal pretreatment. Levels of MEIS2 protein were normalized to Actin and calculated against the untreated control (lane 1). **f** Immunoblotting of indicated proteins in MM.1S and CAG cells treated with Pal for 96 h and Len (3 µM daily) for 72h. Levels of MEIS2 and CRBN proteins were normalized to Actin and calculated against the untreated control. The ratios of MEIS2 to CRBN protein were shown at the bottom. **g** RNA-seq analysis of MM.1S and CAG cells treated with Pal and Len for time indicated.

These data provided the first evidence linking depletion of IKZF1/3 and IRF4 proteins in response to Len and CDK4/6i priming in primary myeloma cells. They further demonstrated that CDK4/6i priming promoted Len signaling by lowering the MEIS2 to CRBN ratio through opposing regulation of MEIS2 and CRBN proteins.

CDK4/6i priming similarly enhanced Len and Pom killing in human myeloma cell lines (HMCL)s MM.1S, CAG in which *TP53* is deleted (Supplementary Fig. 2a) and KMS12PE that harbors a mutated *TP53* ^41^ (Fig. 3b) by caspase-dependent apoptosis (Supplementary Fig. 2b, c). Len slowly induced late G1 arrest in a fraction of MM1.S cells, given the impaired BrdU uptake (Fig. 3c), sustained pRb, elevated p21 ^42^ and p27 proteins, and reduced SKP2 that mediates p27 and p21 degradation ^43^ as well as cyclin A and CDK2 proteins that promote S phase entry at 48 h (Fig. 3d). Cell death followed by 72h (Supplementary Fig. 2d, e). Inhibition of CDK4/6 led to early G1 arrest by 24h regardless of Len, as evidenced by the loss of pSRb and late-G1 core cell cycle proteins (Fig. 3d). This restricted mRNA expression to genes programmed for early G1 only, even in CAG cells that were refractory to cell cycle control by Len (Supplementary Fig. 2a). Confirming its specificity, CDK4/6i failed to induce early G1 arrest or potentiate the reduction of IRF4 in Len killing of U266 cells expressing no Rb, the obligatory CDK4/6 substrate (Supplementary Fig. 2f, h). CDK4/6i priming therefore sensitizes Rb-expressing myeloma cells to IMiD independent of p53 by transcriptional reprogramming and PTR in early G1 arrest.

However, reduction of MEIS2 protein in primary myeloma cells after long exposure to Len or CDK4/6i (Fig. 3a) contrasted the increased in MEIS2 after acute Len treatment in MM1.S cells ^21^. We found that the MEIS2 protein was indeed modestly increased after a brief exposure to Len (1 to 4h) in MM1.S cells, but this increase was obliterated by CDK4/6i priming for 24h (fig. 3e). Moreover, as in MM10, long exposure to Len (72h) or CDK4/6i (96h) profoundly reduced the MEIS2 protein while increasing the CRBN protein cooperatively. This lowered the MEIS2 to CRBN ratio to a level below detection, concurrent with accelerated depletion of IKZF1, IKZF3 and IRF4 and caspase activation in MM1.S cells (Fig. 3f). In CAG cells, CDK4/6i, but not Len, similarly reduced the MEIS2 protein and increased the CRBN protein in cooperation with Len to markedly lower the MEIS2 to CRBN ratio (6-fold) and accelerate IKZF3 (IKZF1 is silenced) and IRF4 depletion and caspase activation (Fig. 3f). There was no change in *CRBN*, *IKZF1* and *IKZF3* mRNAs. However, MEIS2 mRNA and protein were downregulated cooperatively by Len and CDK4/6i priming in MM1.S cells. *IRF4* mRNA was repressed by Len in MM1.S cells as in MM10, and by Len CDK4/6i priming in CAG cells (Fig. 3g), suggesting cell-specific transcriptional and post-transcriptional regulation of MEIS2 and IRF4 by Len and CDK4/6i priming.

CDK4/6 inhibition, therefore, mitigates the rise of MEIS2 protein in acute Len response and inversely regulates MEIS2 and CRBN proteins in cooperation with Len with time to lower the MEIS2 to CRBN ratio that accelerates IKZF3, IKZF1 and IRF4 depletion and caspase activation in myeloma cells.

### CDK4/6i accelerates the displacement of MEIS2 from CRBN to promote IMiD signaling

Our findings further raise the hypothesis that CDK4/6i priming enhances the CRL4^CRBN^ activity by both accelerating the displacement of MEIS2 from CRBN in acute response to Len and destabilizing the MEIS2 protein with time. To test this, we first showed by shRNA knockdown that CRBN was required for Len killing as reported ^8, 9^, more stringently with CDK4/6i priming (Fig. 4a). Next, we demonstrated that at equal loading, the proportion of MEIS2 in the CRBN immunocomplex was reduced to 24% by CDK4/6i priming for 24h, and further to 10% after addition of Len for one hour, in striking contrast to reducing to ∼80% by Len alone in both MM1.S and CAG cells (Fig. 4b). CDK4/6 inhibition therefore rapidly accelerates the displacement of MEIS2 from CRBN by Len in myeloma cells.

**Fig. 4.**
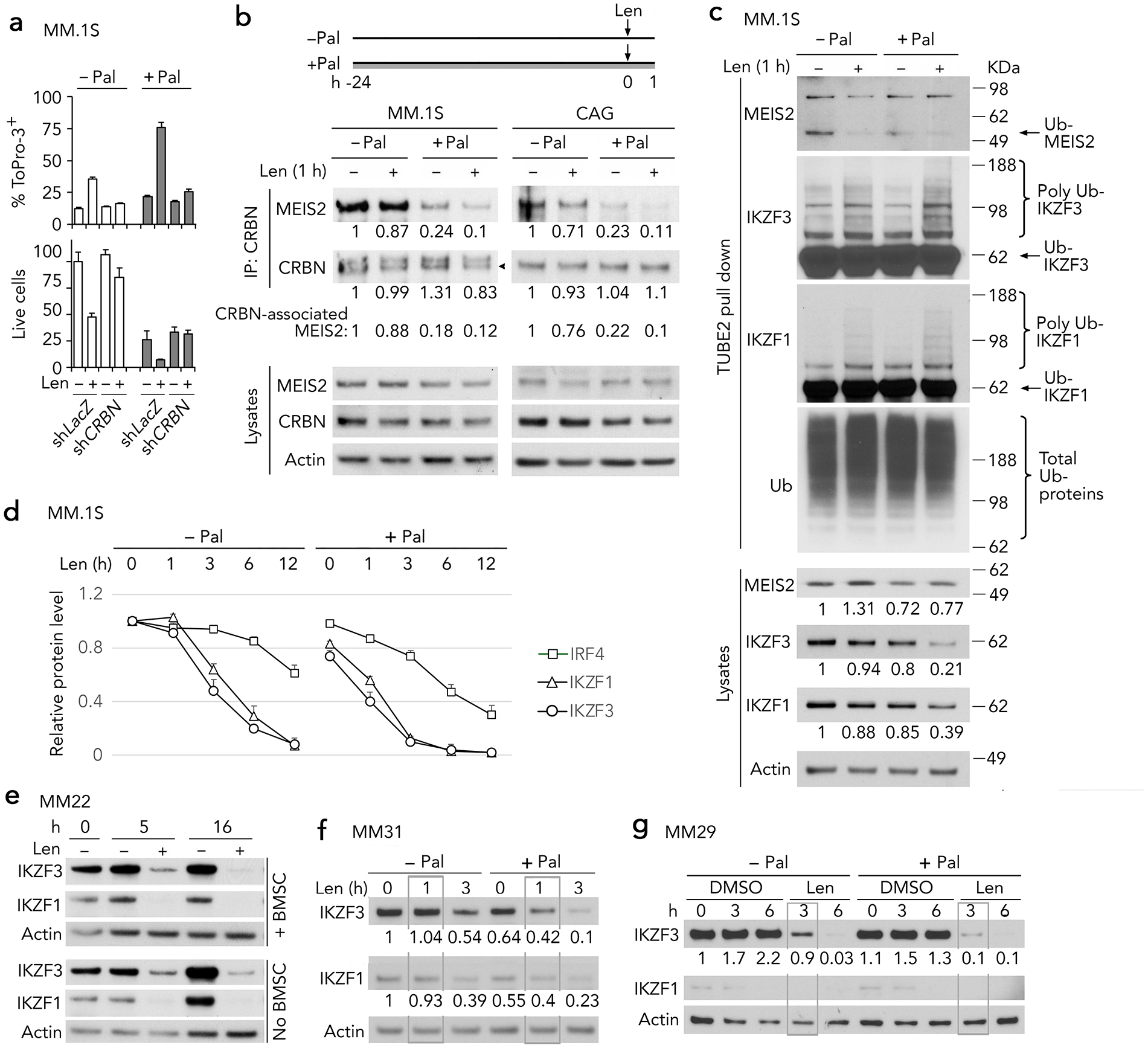
CDK4/6i accelerates displacement of MEIS2 from CRBN to promote IMiD signaling. **a** Percentage of dead (ToPro-3+) and live MM.1S cells stably infected with *LacZ-29 (LacZ)* or *CRBN* shRNA lentivirus and treated with Len (5 μM) daily for 3 days +/- 24 h Pal pretreatment. **b** Immunoprecipitation of CRBN in MM.1S and CAG cells cultured with DMSO or Len for 1 h +/- 24 h Pal pretreatment as diagrammed (top) and immunoblotting of MEIS2 and CRBN in the CRBN immunoprecipitates and the lysates (bottom). The protein level of MEIS2 or CRBN in the lysates was normalized to actin and the relative quantities of MEIS2 and CRBN immunoprecipitated by CRBN antibody (IP) were adjusted based on the normalized CRBN in the lysates. **c** Analysis of ubiquitinated MEIS2, IKZF3 and IKZF1 proteins by TUBE2 pull-down and immunoblotting in MM.1S cells cultured with DMSO or Len as shown in B. Higher molecular weight polyubiquitinated IKZF3 (Poly Ub-IKZF3) and IKZF1 (Poly Ub-IKZF1) are indicated. The levels of MEIS2, IKZF3 and IKZF1 proteins were normalized to Actin and calculated against the untreated control (lane 1). Ub, ubiquitin. **d** Relative level of indicated proteins in MM.1S cells cultured with Len for time indicated +/- Pal pretreatment. **e** Immunoblotting of IKZF3 and IKZF1 in BMMCs (MM22) cultured with Len for time indicated with or without BMSCs. **f-g** Immunoblotting of indicated proteins in BMMCs MM31 and MM29 cultured with Len for time indicated +/- Pal pretreatment for 15 h. The levels of IKZF3 and IKZF1 proteins were normalized to Actin and calculated against the untreated control (lane 1).

Dissociation from CRBN is expected to reduce CRL4^CRBN^ ubiquitination of MEIS2. Indeed, pull-down of ubiquitinated (Ub) proteins with TUBE2 agarose beads revealed that Ub-MEIS2 was reduced by 24h of CDK4/6 inhibition or one hour of Len exposure (Fig. 4c). The total MEIS2 in the lysate was increased by Len as expected but decreased after CDK4/6i priming as we observed (Fig. 3e), despite accelerated dissociation from CRBN (Fig. 4c). Thus, beyond accelerating the displacement of MEIS2 from CRBN by Len, CDK4/6i priming destabilizes the MEIS2 protein in early G1 arrest.

Reduction of Ub-MEIS2 by CDK4/6 priming was reciprocal to increase in Ub-IKZF3 and loss of total IKZF3 in cooperation with acute Len treatment (Fig. 4c). Although changes in Ub-IKZF1 and total IKZF1 were modest initially (Fig. 4c), IKZF1 was depleted in parallel with IKZF3 by 12h of Len treatment, and by 6h with CDK4/6i priming (Fig. 4d and supplementary Fig. 3a). With a delay, the IRF4 protein was reduced gradually in response to Len, consistent with being a direct target of IKZF1/3, and this was accelerated and deepened by CDK4/6i (Fig. 4d and supplementary Fig. 3a). Further confirming its specificity, CDK4/6i did not enhance Len-mediated loss of IKZF3 and IKZF1 in the Rb-null U266 cells (Supplementary Fig. 3b).

Validating the physiologic relevance, IKZF1 and IKZF3 were virtually depleted by 5h of Len treatment in BMMCs (MM22) independent of stromal cells (fig. 4e). CDK4/6i priming accelerated and deepened Len-mediated reduction of IKZF3 and IKZF1 proteins in BMMCs, to complete depletion by 3h in MM31 and 6h in MM29 from patients before achieving PR and VGPR respectively in IMiD therapy (Fig. 4f, g).

Thus, CDK4/6i priming rapidly accelerates the displacement of MEIS2 from CRBN by Len and destabilizes MEIS2 to promote CRL4^CRBN^ ubiquitination-degradation of IKZF1 and IKZF3, leading to downregulation of IRF4 in BMMCs in the context of the clinical response to IMiD.

### CDK4/6i enhances Len killing by activating the IRF4-IRF7-IFN-TRAIL pathway

IRF4 is required for survival of HMCLs ^12^. In separate studies of HMCLs, Len was shown to downregulate IRF4 ^44^ and activate interferon (IFN)-stimulated genes (ISG)s ^45^, and that TRAIL (*TNFSF10*) regulated type I IFN-mediated apoptosis ^46^. Loss of IRF4 relieved repression of IRF7 that mediated the type I IFN response in ABC-DLBCL cell lines ^47^, but the role of IRF7 in Len signaling has not been explored in myeloma cells.

Cooperative regulation of the IRF4 protein by CDK4/6i priming and Len in primary MCL cells and HMCLs (Fig. 3a; Fig 3f, g and Fig. 4d) suggests a pivotal role for IRF4 in orchestrating CDK4/6i priming and proximal Len signaling to apoptosis in myeloma cells. To address this unbiasedly in the context of clinical response to IMiDs, we analyzed BMMCs from myeloma patients before achieving CR (MM18) or VGPR (MM10, 11 and 19) ( Group I) and refractory patients (MM 20, MM30) (Group II) in IMiD therapy (Fig. 2f-h and supplementary Table 1) by RNA-seq after 24h of Len exposure with or without CDK4/6i priming *ex vivo*.

The number of Len-regulated genes in Group I BMMCs was markedly higher compared with Group II BMMCs and amplified by CDK4/6i priming (Fig. 5a and supplementary Table 3). Gene set enrichment analysis of Group I vs Group II BMMCs further revealed a significant enrichment of the IFN response gene signatures by Len and by CDK4/6i (Pal) (Nominal p-value < 0.01), greater when combined (Nominal p-value < 0.0001) (Fig. 5b and supplementary Table 4). In particular, the type I IFN response genes such as *DDX60*, *IFI44L, XAF1, MX1, TNFSF10, and IRF7* were differentially upregulated by Len in individual Group I BMMCs, greater with CDK4/6i priming and correlating with Len killing (Fig. 2f and Fig. 5c).

**Fig. 5.**
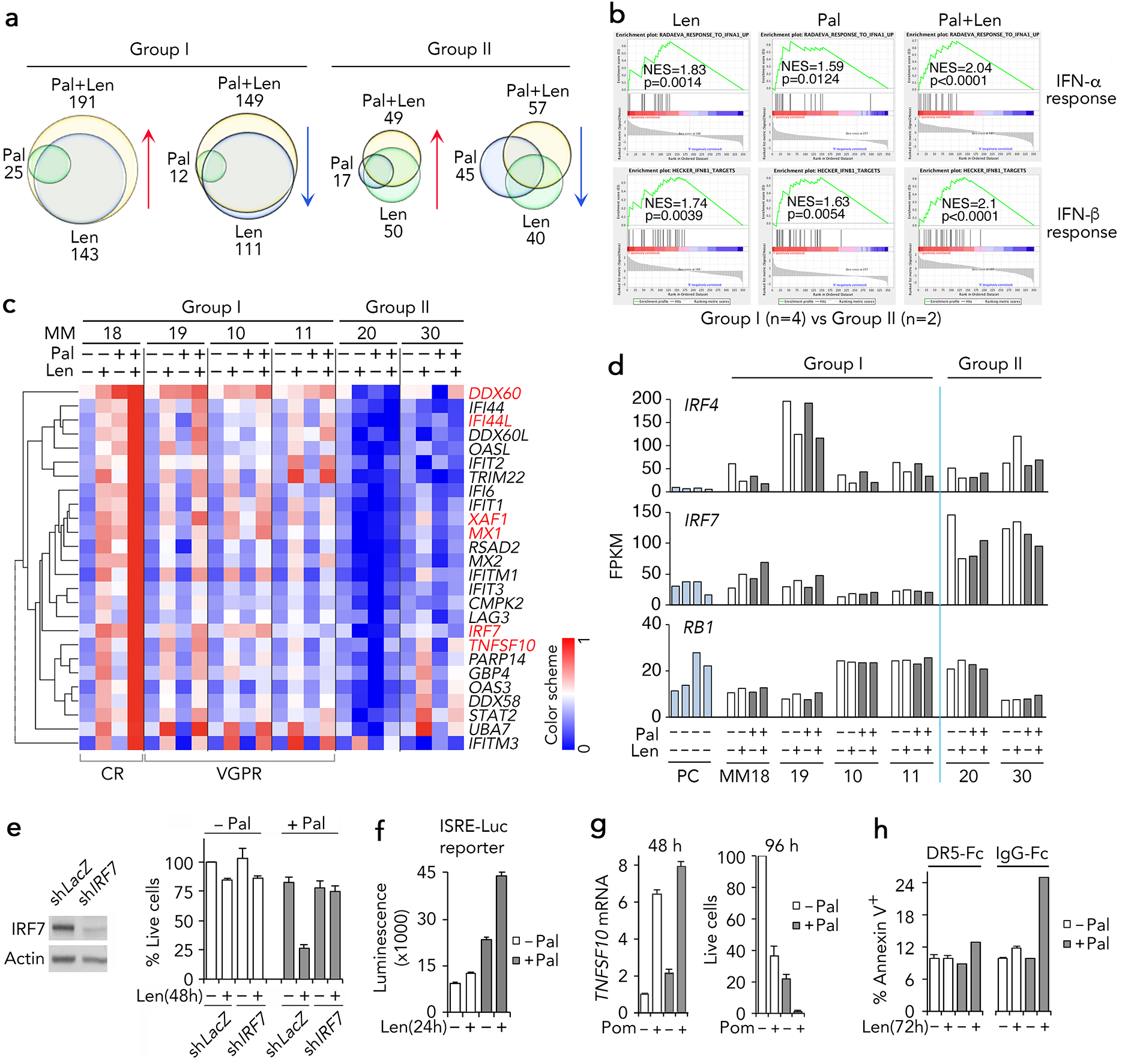
CDK4/6i enhances Len killing by activating the IRF4-IRF7-IFN-TRAIL pathway. **a** Venn diagrams of differentially expressed genes indicated by RNA-Seq analysis of freshly isolated BMMCs treated with Len (Len, 3 µM) with or without Pal pretreatment for 6 h or Pal (Pal, 300 nM) only for 24 h. Group I refers to BMMCs from patients who responded to the first post-biopsy Len-based therapy and Group II from patients who were resistant to Len-based therapy before biopsy. **b** GSEA identified enrichment of the IFN responsive gene sets that distinguish Group I from Group II BMMCs. **c** Heatmap of IFN signature genes in BMMCs treated as in (**a**). **d** RNA-Seq analysis of mRNA abundance of indicated genes in PCs and BMMCs treated as in (**a**). **e** Immunoblotting of IRF7 in MM.1S cells stably infected with the *LacZ-29* shRNA (sh*LacZ*) or *IRF7* shRNA (sh*IRF7*) lentivirus (left), and cell viability after exposure to Len (10 μM) for 48 h +/- Pal pretreatment (right). **f** Luciferase activity measured by luminiscence in MM.1S cells carrying an inducible ISRE-responsive luciferase reporter construct (ISRE-Luc reporter) treated with 3 μM Len for 24 h +/- Pal-pretreatment. **g** q-PCR analysis of *TNFSF10* in MM.1S cells exposed to Pom (3μM) for 48 h +/-Pal pretreatment (left). The percentage of live MM.1S cells was determined at 96 h (right). (H) Analysis of Annexin V^+^ MM.1S cells cultured with Len for 72 h +/- Pal pretreatment and DR5-Fc or IgG-Fc was added 1 h before Len on days 0 and 2. Data are representative of three independent experiments.

Of interest, *IRF4* mRNA was downregulated by Len in Group I BMMCs, with reciprocal increase in *IRF7* mRNA in BMMCs from patients before CR (MM18) and VGPR (MM19) (Fig. 5d). IRF4 antagonizes CDK4/6i priming, as *IRF4* knockdown exacerbated Len killing of KMS12-PE cells with CDK4/6i priming to near extinction (Supplementary Fig. 4a). Conversely, IRF7 was required for CDK4/6i priming, because knocking down *IRF7* abrogated the enhancement of marginal Len killing at 48h by CDK4/6i priming in MM.1S cells (Fig. 5e). Len and CDK4/6i priming cooperatively upregulated *IRF7* and *IFNB* (Supplementary Fig. 4b) and activated a luciferase reporter carrying an ISG element (Fig. 5f). Accordingly, exogenous IFNβ enhanced Len killing and CDK4/6i priming (Supplemental Fig. 4c). Mimicking the prominent upregulation of *TNFSF10* mRNA by Len and CDK4/6i priming in MM18 (Fig. 5c), *TNFSF10* mRNA was profoundly upregulated by Pom and CDK4/6i priming, leading to virtual depletion of MM.1S cells (Fig. 5g). Lastly, blocking TRAIL with DR5-Fc, but not with the control IgG-Fc ^48^, rescued MM.1S cells from Len killing with CDK4/6i priming (Fig. 5h), further demonstrating that TRAIL mediates IMiD killing and CDK4/6i priming.

Collectively, our data demonstrated, for the first time, that through opposing regulation of MEIS2 and CRBN proteins CDK4/6i priming promotes proximal Len signaling to downregulate IRF4, de-repress IRF7 and enhance the IFN response and TRAIL-mediated killing of primary myeloma cells in the context of clinical response to IMiDs.

### CDK4/6i enhances CELMoD killing by lowering the MEIS2 to CRBN ratio that promotes IRF4-IRF7-IFN-TRAIL signaling

The CELMoD iberdomide (Ibd) is more potent than Len or Pom in myeloma therapy owing to its high affinity for CRBN ^13, 14^. Ibd killing of the *TP53*-null CAG cells at the saturating dose (100 nM) was comparable to Len (3 µM) and enhanced by CDK4/6i priming over a 100-fold range (Fig. 6a). Like Len, Ibd did not regulate the cell cycle in CAG cells given the sustained TK1 expression, or affect induction of early G1 arrest by CDK4/6i as indicated by profound repression of *TK1* and *AURKB* mRNAs within 24h (Fig. 6b). Ibd killing of myeloma cells is therefore significantly more effective than Len but also enhanced by CDK4/6i priming independent of p53.

**Fig. 6.**
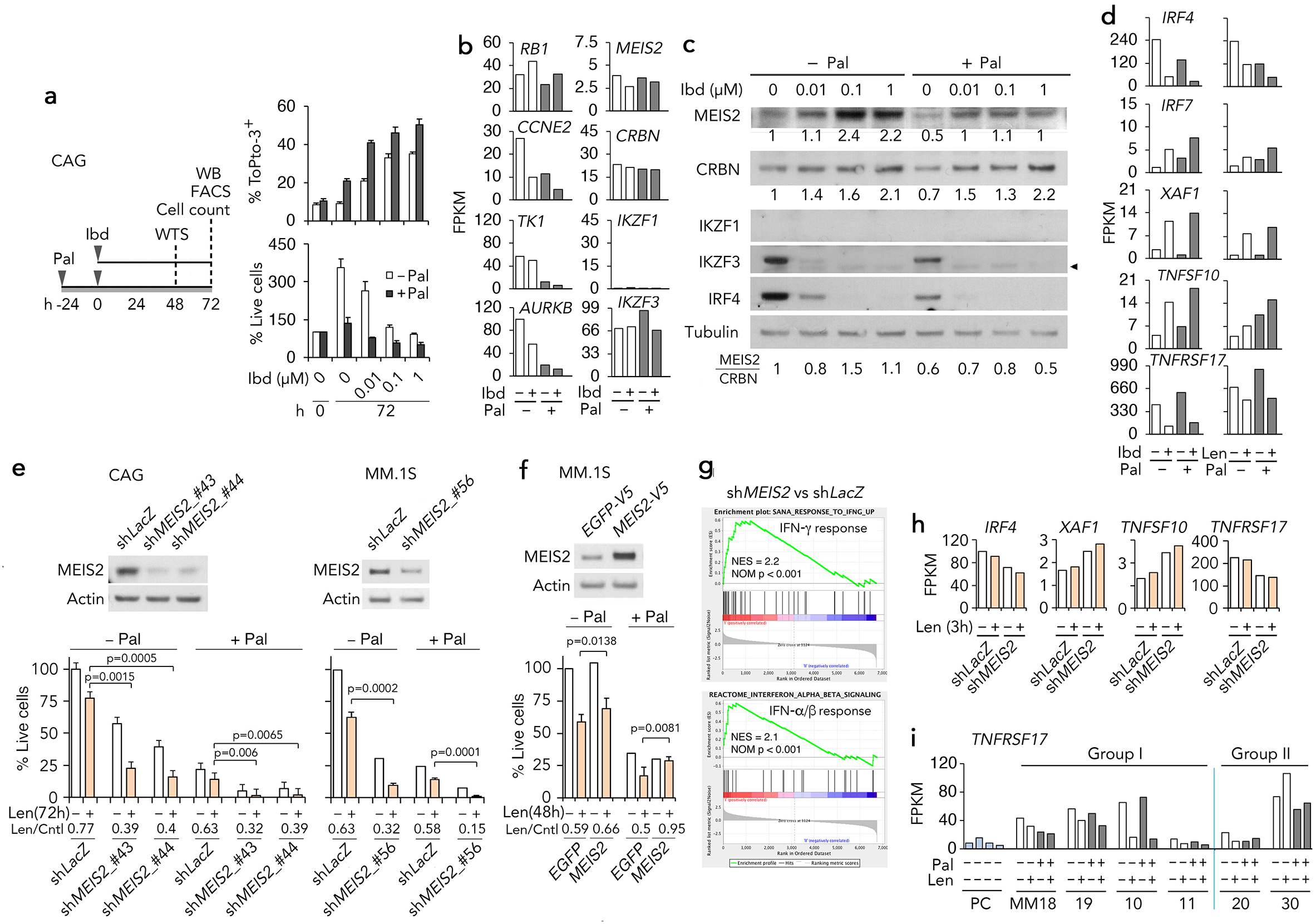
CDK4/6i enhances CELMoD killing by reducing MEIS2 which sustains BCMA expression and antagonizes its repression by CELMoD and IMiD. **a** Outline for Pal and Iberdomide (Ibd) treatment and time of analysis (left), the percentage of dead cells (ToPro-3+) and live cells relative to input at 72 h of culturing with Ibd at indicated concentrations with Pal pretreatment (right). **b** RNA-seq analysis of mRNA abundance of indicated genes in CAG cells treated with Ibd (100 nM) for 48 h with or without Pal pretreatment. **c** Immunoblotting of indicated proteins in CAG cells treated with Ibd at indicated concentrations for 72 h with or without Pal pretreatment. The protein level of MEIS2 or CRBN was normalized to Tubulin and calculated against the untreated control (lane 1). The ratios of MEIS2 to CRBN protein were shown at the bottom. **d** RNA-seq analysis of mRNA abundance of indicated genes in CAG cells treated with Ibd (100 nM) for 48 h or Len (3 µM) for 72 h with or without Pal pretreatment. **e** Immunoblotting of MEIS2 in CAG and MM.1S cells infected with *LacZ-29* shRNA (sh*LacZ)*, or sh*MEIS2*_#43 (TRCN0000016043), sh*MEIS*2_#44 (TRCN0000016044) or shMEIS2_#56 (TRCN0000274056) lentivirus. Cell viability was determined after 72 h of Len exposure (5 μM) +/- Pal pretreatment (bottom). **f** Immunoblotting of MEIS2 in MM.1S cells stably infected with pLX304*-EGFP (V5-EGFP)* or *MEIS2* (*V5-MEIS2*) overexpressing lentivirus (top). Cell viability was determined after 96 h of Len exposure (10 μM) +/- Pal pretreatment (bottom). **g-h** Enrichment of IFN responsive gene sets (**g**) and expression of indicated genes (**h**) in MM.1S cells infected with *MEIS2* shRNA (sh*MEIS2*) or sh*LacZ*. **i** RNA-Seq analysis of mRNA abundance of *TNFRSF17* in PCs and BMMCs treated as in Figure 5A.

MEIS2 and CRBN were increased in a dose-dependent manner after long exposure (72h) to Ibd in CAG cells (Fig. 6c). The increase in MEIS2 protein was likely due to stabilization after it was efficiently displaced from CRBN by the high affinity Ibd. CDK4/6i priming reduced the MEIS2 protein in all myeloma cells (Fig. 3a, e and Fig. 6c), and blunted Ibd-mediated rise of MEIS2 protein while increasing the CRBN protein. This resulted in lowering the MEIS2 to CRBN ratio, which correlated with enhanced Ibd killing by CDK4/6i priming (Fig. 6c). In addition, Ibd downregulated the *IRF4* mRNA, and inversely upregulated *IRF7, XAF1* and *TNFSF10* mRNAs more prominently than Len at 1/30 concentration, and this was also enhanced by CDK4/6i priming (Fig. 6d). Altogether, these results suggest that the IRF4-IRF7-IFN-TRAIL pathway mediates Ibd’s robust anti-myeloma activity, and this was enhanced by CDK4/6i priming that lowers the MEIS2 to CRBN ratio to mitigate MEIS2 inhibition of CRL4^CRBN^.

### MEIS2 sustains BCMA expression and antagonizes repression of BCMA by Ibd and Len

Of interest, *TNFRSF17* mRNA encoding BCMA for survival of myeloma cells ^49, 50^ was more prominently repressed by Ibd than by Len, but not by CDK4/6i priming (Fig. 6d), invoking a previously unrecognized mechanism of Ibd and IMiD killing in myeloma cells. To address this and confirm that MEIS2 inhibits IMiD killing and CDK4/6i priming in myeloma cells, we knocked down MEIS2 in two HMCLs using 3 shRNA clones. Knocking down MEIS2 was toxic to MM.1S cells as reported ^51^ as well as CAG cells, and exacerbated Len killing with CDK4/6i priming in both HMCLs (Fig. 6e and supplementary Fig. 5a). Confirming the MEIS2 specificity, ectopic expression of MEIS2 attenuated Len killing and rescued MM1.S cells from Len killing with CDK4/6i priming (Fig. 6f). Moreover, MEIS2 knockdown downregulated the *IRF4* mRNA, enriched the IFN response signature (Supplemental Fig. 5b) and upregulated *XAF1* and *TNFSF10* mRNAs in MM.1S cells (Fig. 6h), which mimics the enhancement of Ibd and IMiD killing by CDK4/6i priming in HMCLs and BMMCs (Fig. 3b, d and Fig. 6d). MEIS2 therefore antagonizes IRF4-IRF7-IFN-signaling to TRAIL-mediated apoptosis downstream of CRL4^CRBN^ in response to IMiD and CDK4/6i priming in myeloma cells.

Knocking down MEIS2 downregulated *TNFRSF17* mRNA with or without Len treatment, revealing a requirement for MEIS2 in sustaining BCMA expression for intrinsic survival of myeloma cells and for repression of *TNFRSF17* by Len (Fig. 6h). Importantly, *TNFRSF17* mRNA was downregulated by Len and not by CDK4/6i priming in BMMCs from MM patients responding (Group I) but not refractory (Group II) to IMiD therapy (Fig. 6i). IMiD and Ibd kill myeloma cells by both repressing the BCMA survival pathway and activating the IRF4-IRF7-IFN-TRAIL pathway. MEIS2 antagonizes IMiD and Ibd therapy by both inhibiting CRL4^CRBN^ and sustaining BCMA in the context of the clinical response to IMiD therapy.

### Mechanism of CDK4/6i reprogramming for IMiD and CELMoD vulnerability in myeloma

Our data have uncovered that MEIS2 inhibits the CRL4^CRBN^ activity and sustains BCMA expression for survival of myeloma cells, and a novel mechanism by which CDK4/6 inhibition reprograms myeloma cells for IMiD and CELMoD vulnerability (Fig. 7). **A**. MEIS2 binds CRBN and inhibits the CRL4^CRBN^ activity. Expression of BCMA and activation of IRF4 by IKZF3 and IKZF1 and other transcription factors maintain the survival of myeloma cells. **B**. IMiD (Len) or Ibd displaces MEIS2 from CRBN and recruit IKZF3 and IKZF1 to CRL4^CRBN^ for ubiquitination and degradation. This, in turn, downregulates IRF4 to relieve IRF7 for induction of IFN and TRAIL-mediated apoptosis. Len also reduces the MEIS protein required for BCMA expression. **C**. CDK4/6 Inhibition induces early G1 arrest that disrupts the association of MEIS2 with CRBN and destabilizes the MEIS2 protein while increasing the CRBN protein with time to lower the MEIS2/CRBN ratio. **D**. CDK4/6i priming rapidly accelerates the displacement of MEIS2 from CRBN by Len, leading to depletion of IKZF1/3 and IRF4 that promotes TRAIL-mediated apoptosis in concert with loss of IRF4 by CDK4/6i priming and BCMA expression in response to Len and Ibd. By lowering the MEIS2 to CRBN ratio through opposing regulation of MEIS2 (purple bar) and CRBN proteins with time (yellow bar), CDK4/6i priming mitigates MEIS2 inhibition of CRL4^CRBN^ in Len and Ibd signaling and kills myeloma cells in concert with reduction of BCMA by Len (Ibd).

**Fig. 7.**
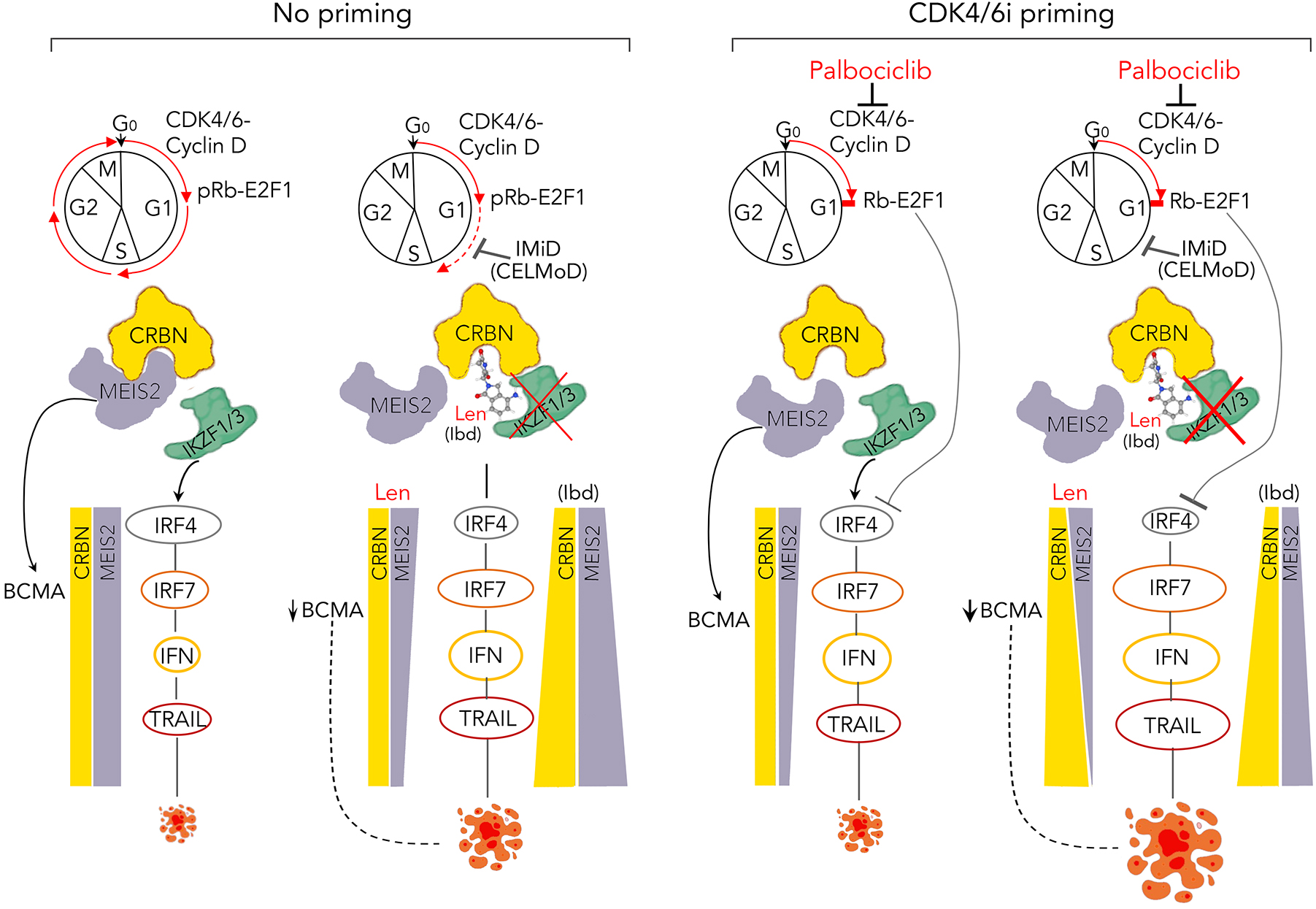
Mechanism for CDK4/6i reprograming of myeloma cells for IMiDs and CELMods vulnerability. Binding of lenalidomide (Len) or Iberdomide (Ibd) to CRBN displaces the endogenous substrate MEIS2 from CRBN within one hour. This promotes the recruitment of IKZF1 and IKZF3 to CRL4^CRBN^ for ubiquitination and degradation (Proximal), and loss of the IKZF1/3 target IRF4 that leads to induction of interferon (IFN) and TRAIL-mediated apoptosis (Distal). Inhibition of CDK4/6 by palbociclib for 24h also displaces MEIS2 from CRBN and enhances apoptosis via the IRF4-IFN-TRAIL pathway independent of Len, and cooperatively with Len recruitment of IKZF1/3. With time, Pal and Len cooperatively repress MEIS2 and increase CRBN proteins, leading to enhanced apoptosis. IFN, interferon; BCMA, protein encoded by *TNFRSF17*.

## Discussion

We have discovered that MEIS2 functions as an inhibitor of CRL4^CRBN^ to promote resistance to IMiD and CELMoD in multiple myeloma. First, expression of high *MEIS2, CDK4, CDK6* and low *CRBN* in BMMCs before therapy strongly correlated with inferior OS in IMiD monotherapy in the CoMMpass clinical trial (Fig. 1). Second, IMiD and CELMoD exert their anti-myeloma activity mainly by inducing TRAIL-mediated apoptosis downstream of the CRBN-IKZF1/3-IRF4-IRF7-IFN signaling pathway (Fig. 3; Fig. 5 and Fig. 6), and MEIS2 inhibits this pathway in Len response and CDK4/6i priming (Fig. 6). Third, by lowering the MEIS2 to CRBN ratio, CDK4/6i priming enhances IMiD and Ibd killing of HMCLs (Fig. 3 and Fig. 6) and more importantly, IMiD killing of BMMCs *ex vivo* in the context of the clinical response to IMiD *in vivo* (Fig. 2; Fig. 3 and Fig. 4).

Furthermore, we’ve discovered that MEIS2 sustains *TNFRSF17* expression, which explains why knocking down MEIS2 was toxic to myeloma cells; that correlating with more effective killing, Ibd more prominently represses *TNFRSF17* than Len (Fig. 6). These data strongly suggest that MEIS2 promotes survival of myeloma cells by maintaining BCMA (encoded by *TNFRSF17*) signaling. *TNFRSF17* was downregulated by Len in BMMCs from IMiD-responding but not refractory patients (Fig. 6) further supporting its clinical relevance. MEIS2 is a transcription factor required for limb, brain, lens and retina development in mice ^15, 52, 53, 54, 55, 56, 57, 58^ and acts as an oncogenic partner for ETO-AML in a mouse model of AML ^20^. Overexpression of MEIS2 has been associated with proliferation and survival in neuroblastoma and ovarian cancer cell lines ^59, 60^. That MEIS2 both inhibits CRL4^CRBN^ activity to antagonize response to IMiD and Ibd and sustains BCMA expression to promote survival of myeloma cells suggest MEIS2 as a tumor promoter in myeloma. MEIS2 is aberrantly expressed in myeloma cells and not in normal B cells or other malignant B cells (Fig. 1; Di Liberto, Huang and Chen-Kiang, unpublished), further providing a rationale for enhancing IMiD and CELMoD therapy by targeting MEIS2.

CDK4 and CDK6 are dysregulated at a high frequency in cancer development and progression, including multiple myeloma ^22^. Here we demonstrate for the first time that CDK4/6i priming promotes Len signaling to depletion of IKZF1/3 and IRF4 that relieve IRF7 to enhance the IFN response and TRAIL-mediated apoptosis in primary myeloma cells *ex vivo* in the context of clinical response to IMiDs in patients (Fig. 3 and Fig. 5). Our integrated analysis further revealed a novel bipartite mechanism by which CDK4/6i primes myeloma cells for vulnerability to IMiD by both accelerating the displacement of MEIS2 from CRBN by Len and destabilizing MEIS2 protein in opposition to increasing the CRBN protein with time in early G1 arrest (detailed in Fig. 7). In addition, MEIS2 is downregulated transcriptionally by Len and Ibd and by CDK4/6i in a cell-specific manner (Fig. 3 and Fig. 6). The mechanisms for regulation of MEIS2 and CRBN, such as autoregulation of components of E3 ligases ^61^, remain important questions to follow.

Apart from enhancing the myeloma cell intrinsic sensitivity to IMiD and Ibd (Fig. 2; Fig. 3 and Fig. 6), CDK4/6 inhibition has been reported to reprogram cancer-immune cell interaction in various preclinical solid cancer models; it triggers anti-tumor immunity ^62^, augments anti-tumor immunity by enhancing T-cell activation ^63^, elicits a cancer cell program that enhances the checkpoint blockade therapy ^64^; and promotes tumor immunity through induction of T cell memory ^65, 66^. Various CDK4/6 inhibitors are now FDA-approved for treatment of breast cancer. New and specific agents such as venetoclax ^67^ are available as partners. We established the feasibility of inhibiting CDK4/6 in combination therapy in human cancer in a proof-of concept clinical trial combining palbociclib with bortezomib in multiple myeloma ^68^. We now show that inhibition of CDK4/6 leads to early G1 arrest despite *TP53* deletion or mutations in myeloma cells to enhance the anti-myeloma activity of IMID and CELMoD (Fig. 3 and Fig. 6).

Targeting myeloma-immune cell interaction with BCMA-based bispecific antibody and CAR T-cell therapy is clinically effective. Our discovery that BCMA is repressed by Len and Ibd calls attention to combining IMiD or CELMod with BCMA-based immunotherapy. In contrast, BCMA is not repressed by CDK4/6i priming, despite reduction of MEIS2. Enhancing the clinical efficacy of IMiD and CELMoD by CDK4/6 inhibition, especially in the relapsed and refractory setting, is a rational and exciting prospect.

## Methods

### Isolation of human BMMCs and culture

Bone marrow specimens were obtained from multiple myeloma patients at New York-Presbyterian Hospital under informed consent as part of an Institutional Review Board approved study. Primary CD138^+^ human BM myeloma (BMM) cells were isolated as previously described ^24^ and co-cultured at a 4:1 ratio, with mitomycin C-arrested HS-5 stromal cells (RRID:CVCL_3720, kindly provided by Dr. Hazlehurst at Lee Moffitt Cancer Center Tampa, FL) in the presence of recombinant human IL-6 (40 U/ml, R&D Systems, 206-IL), soluble IL-6 receptor (40 U/ml, R&D Systems, 8954-SR) and human IGF-1 (100 ng/ml, R&D Systems, 291-G1)^25^. For analysis, BMMCs were removed from stromal cells by gently pipetting.

### Gene expression and patient overall survival analysis

Gene expression and clinical data of MMRF CoMMpass study (https://clinicaltrial.gov entry NCT01454297) were obtained from The Cancer Genome Atlas (TCGA) using the Genomic Data Commons (GDC) portal ^29^. Myeloma patients from the CoMMpass study were at least 18 years old including both male and female sex. Baseline gene expression counts in Fragments Per Kilobase of exon model per Million mapped reads (FPKM) were downloaded for all samples. Clinical data of overall survival time were retrieved and matched to expression data. Patients were divided into “High” and “Low” expression groups using indicated gene expression value as the cutoff. Overall survival curves were estimated using the Kaplan–Meier method with the survival package in R (version 4.4.2, RRID:SCR_001905). Group differences were assessed using the log-rank test ^30, 31^.

### MM cell lines and culture

MM.1S (RRID: CVCL_8792) and CAG (RRID: CVCL_D569) human myeloma cell lines (HMCL) were kindly provided by Dr. N. Krett (Northwestern University, Chicago, IL), and KMS-12-PE (RRID: CVCL_1333) by Dr. Louis Staudt (NIH). Rb-deficient U266 (RRID: CVCL_0566) cells were purchased from ATCC (Rockville, MD). HMCLs were cultured as described previously^25^. No authentication of these cell lines was done by the authors.

### Drugs and treatment

Unless otherwise indicated, palbociclib (Selleck Chemicals, S1116) was added to HMCLs (0.3 μmole/L for 24 hours) and to primary BMMCs (0.5 μmole/L for 5 hours) before addition of lenalidomide (Selleck Chemicals, S1029), pomalidomide (Selleck Chemicals, S1567) (3 μmoles/L to HMCLs and 5 μmoles/L to BMMCs) or iberdomide (0.01 to 1 μmoles/L, Selleck Chemicals, S8760). Pan caspases inhibitor Q-VD-OPh (20 μmoles/L, BD Biosciences, OPH001, RRID:AB_2869523), human DR5-Fc (100 ng/ml, ACROBiosystems, TR2-H5255-100UG), or Fc portion of human IgG (100 ng/ml, Rockland, 609-1103) was added 1 hour before addition of lenalidomide or pomalidomide as indicated.

### Cell cycle and cell death assays

Analysis of the cell cycle by 5-bromo-2-deoxyuridine (BrdU) (Sigma-Aldrich, B5002) uptake in 30 minutes and DNA content per cell by propidium iodide staining, cell death assays by ToPro-3 (Thermo Fisher Scientific, T3605) or MitoTracker Red (Thermo Fisher Scientific, M46752) staining, or by using eBioscience^TM^ Annexin V Apoptosis Detection Kit (Thermo Fisher Scientific, 88-8005-74), and determination of total live cells by trypan blue exclusion were performed as described previously ^25^.

### RNA-Seq and analysis

Total RNA was isolated using the RNAEasy kit (QIAGEN, 74104). 100 ng of RNAs were converted to cDNA, which was isolated with magnetic beads from the TruSeq mRNA prep kit (v2) (Illumina, RS-122-2102) and then ligated to Illumina adapters. Using these multi-plexed cDNA libraries, we generated clusters on the Illumina cBot station and paired-end sequenced each sample to 50x50 bp on the Illumina HiSeq2000 Sequencing System (RRID:SCR_020130). Cluster generation, sequencing, and processing of the images were done on the HiSeq2000 and post-processing of reads was performed with CASAVA (v.1.8.2) (RRID:SCR_001802). Raw sequencing reads with a minimum of about 50 million reads per sample were cleaned by trimming adapter sequences and low quality bases using cutadapt (RRID:SCR_011841) ^32^, and were aligned to the human reference genome (GRCh37) using STAR (RRID:SCR_004463) ^33^. Cufflinks (RRID:SCR_014597) was used to measure transcript abundances in FPKM ^34, 35^. Heatmap of gene expression with genes clustered using the Euclidean algorithm was generated by using Morpheus (RRID:SCR_014975) (https://software.broadinstitute.org/morpheus/).

### Gene set enrichment analysis

Gene Set Enrichment Analysis (RRID:SCR_003199) ^36^ was performed using C2 curated functional gene sets in the Molecular Signature Database (MSigDB). The normalized enrichment score (NES) is used to compare gene set enrichment across gene sets and the nominal p value to estimate the statistical significance of the enrichment score for a single gene set. Gene sets having nominal *p* values no more than 0.05 were considered significant in enrichment.

### Quantitative RT-PCR

The first strand cDNA was synthesized from total RNA using SuperScript III (Thermo Fisher Scientific, 18080400) and subjected to real-time RT-PCR using Assays-on-Demand gene expression mixes specific for each human gene with *ACTB* as control and the TaqMan Universal PCR Master Mix (Thermo Fisher Scientific, 4304437). Reactions were carried out in triplicate in the Life Technologies QuantStudio 6 Real Time PCR System (RRID:SCR_020239). The relative amount of product was determined by the comparative Ct method.

### Immunoblotting

Immunoblotting was performed as described ^25^. Antibodies against phosphorylated Rb (Ser807/811)(9308, RRID:AB_331472), Rb (9309, RRID:AB_823629), CDK4 (12790, RRID:AB_2631166), Cyclin D1 (2922, RRID:AB_2228523), Cyclin D2 (3741, RRID:AB_2070685), Cyclin D3 (2936, RRID:AB_2070801), cleaved caspase-3 (9661, RRID:AB_2341188), and IKZF1 (5443, RRID:AB_10691693) were purchased from Cell Signaling Technology; actin (612656, RRID:AB_2289199) from BD Biosciences; CDK6 (sc-7961, RRID:AB_627242), cyclin A (sc-751, RRID:AB_631329), cyclin E (sc-481, RRID:AB_2275345), p21 (sc-397, RRID:AB_632126), p27 (sc-528, RRID:AB_632129), SKP2 (sc-7164, RRID:AB_2187650), IRF4 (sc-48338, RRID:AB_627828) and IRF7 (sc-9083, RRID:AB_2127436) from Santa Cruz; IKZF3 (IMG-6283A, RRID:AB_1930192) from IMGENEX, MEIS2 (HPA003256, RRID:AB_1079356) from Sigma-Aldrich; and CRBN (NBP1-91810, RRID:AB_11037820) from Novus Biologicals. Peroxidase AffiniPure Goat anti-Mouse IgG (115-035-062, RRID: AB_2338504) and Peroxidase AffiniPure Donkey anti-Rabbit IgG (711-035-152, RRID: AB_10015282) were purchased from Jackson ImmunoResearch.

### shRNA knockdown

The *IRF4* shRNA (sh*IRF4*) (TRCN0000014767), sh*CRBN* (TRCN0000141985), sh*IRF7* (TRCN0000014858), sh*GFP*437, or sh*LacZ-29* (The RNAi Consortium at Broad Institute) lentiviruses were produced as previously described^25^. sh*MEIS2*_#56 (TRCN0000274056), sh*MEIS2*_#43 (TRCN0000016043) and sh*MEIS2*_#44 (TRCN000001604) was purchased from Sigma. Knockdown of each target was validated by quantitative RT-PCR and/or immunoblotting. ***Immunoprecipitation*** MM.1S or CAG cells were treated with DMSO or 3 μM lenalidomide for 1 hour and lysed in Pierce™ IP lysis buffer (Thermo Fisher Scientific, PI87788) containing 1x protease inhibitor cocktails (Sigma-Aldrich, 11836170001) and 10 μM MG132 (Sigma-Aldrich, M7449). CRBN was immunoprecipitated by incubating 2 μg rabbit anti-CRBN antibody (Novus Biologicals, Cat# NBP1-91810, RRID:AB_11037820) with 500 μg protein lysates in 500 μl Pierce™ IP lysis buffer overnight at 4°C. The immunoprecipitates were then incubated with 25 μl Protein G PLUS-agarose beads (Santa Cruz Biotechnology, sc-2003, RRID:AB_10201400) for 1 hour at 4°C and the beads were washed 4 times with IP lysis buffer. Eluted proteins were analyzed by immunoblotting.

### In vivo ubiquitination

MM.1S cells were treated with DMSO or 3 μM lenalidomide for 1 hour and then lysed in Pierce™ IP lysis buffer containing 1x protease inhibitor cocktails, 10 mM NEM (Sigma-Aldrich, E3876) and 10 μM MG132. Ubiquitinated IKZF1 and IKZF3 proteins were pulled down by anti-Ub TUBE2 agarose beads (LifeSensors, UM402) overnight at 4°C, washed 3 times with IP lysis buffer and analyzed by immunoblotting. Total ubiquitinated proteins were detected by using anti-ubiquitin antibody (Enzo Life Sciences, Cat# LSI-AB-0120, RRID:AB_311908).

### Ectopic expression of human MEIS2

pLX304-*MEIS2* (DNASU plasmid repository, RRID:SCR_012185) or pLX304-*EGFP* (Broad Institute, RRID:SCR_007073) *vector* was co-transfected with psPAX2 (RRID:Addgene_12260) and pMD2.G (RRID:Addgene_12259) plasmids into 293T cells (ATCC, CRL-3216, RRID:CVCL_0063) and the generated lentivirus were used to infect MM.1S cells. After selection with Blasticidin S (8 µg/mL) (Thermo Fisher Scientific, A1113903), the cells were analyzed by immunoblotting to confirm the expression of MEIS2.

### ISRE reporter assay

MM.1S cells were transduced with ISRE luciferase reporter lentivirus (SA Biosciences, 79824) and selected with puromycin (Thermo Fisher Scientific, A1113803). The selected cells were treated with lenalidomide (3 μM) for 24 hours with or without palbociclib-pretreatment, and luciferase activity was measured using One-Step^TM^ Luciferase Assay System (SA Biosciences, 60690-2) on a Promega GloMax 20/20 Luminometer (RRID:SCR_018613).

### Statistical analysis

To assess the synergistic effect of two-drug combination, two-way ANOVA was performed on the square root transformed data. A significant interaction from the two-way ANOVA analysis is indication of a synergistic effect. An additive effect is assumed if the interaction is not significant, while each drug treatment alone is significantly different from the combination treatment. Tukey’s Studentized Range (HSD) Test was used to control the Type I familywise error rate when pairwise comparisons were conducted. All tests were two-tailed with p-value 0.05 or less to declare statistical significance. Each experiment included at least three biological replicates per group. A power analysis was not performed prior to each experiment due to the exploratory nature of this study. Whenever conditions permitted, investigators responsible for each experiment and data analysis were blinded to the experimental group assignments. The statistical analyses were done in Statistical Analysis System (RRID:SCR_008567) SAS9.3. Unless otherwise noted, error bars in figures are displayed as ±SEM.

### Data Availability

The data generated in this study are publicly available via NCBI Sequencing Reads Archive (SRA) under accession number PRJNA575377.

## Acknowledgements

This study was supported in part by the STARR Cancer Consortium Grant I3-A162 (S.C-K.) and National Cancer Institute Grants R01 CA188794 (S.C-K.) and R01CA276053 (G.L.). We thank Suzan Chen for technical assistance, and Jenny Zhang and Tuo Zhang at Genomics Resources Core Facility of Weill Cornell Medicine for RNA-sequencing.

## Author Contributions

X.H., M. D. L., R.N., S. C-K. contributed to conception, design, development of methodology, writing manuscript, and study supervision; X.H., M.D-L., T.M.M., G.L., R.N., S.C-K. contributed to analysis and interpretation of data; and X.H., D.J., M.D.L., Z.C., S.E., T.M.M., G.L., R.N., S.C-K. contributed to administrative, technical or material support.

## References

1. Chari A, et al. Daratumumab plus pomalidomide and dexamethasone in relapsed and/or refractory multiple myeloma. Blood 130, 974–981 (2017).

2. Usmani SZ, et al. Pembrolizumab plus lenalidomide and dexamethasone for patients with treatment-naive multiple myeloma (KEYNOTE-185): a randomised, open-label, phase 3 trial. Lancet Haematol 6, e448–e458 (2019).

3. Dimopoulos MA, et al. Addition of elotuzumab to lenalidomide and dexamethasone for patients with newly diagnosed, transplantation ineligible multiple myeloma (ELOQUENT-1): an open-label, multicentre, randomised, phase 3 trial. Lancet Haematol 9, e403–e414 (2022).

4. Dimopoulos MA, et al. Elotuzumab, lenalidomide, and dexamethasone in RRMM: final overall survival results from the phase 3 randomized ELOQUENT-2 study. Blood Cancer J 10, 91 (2020).

5. Sonneveld P, et al. Daratumumab, Bortezomib, Lenalidomide, and Dexamethasone for Multiple Myeloma. N Engl J Med 390, 301–313 (2024).

6. Ito T, et al. Identification of a primary target of thalidomide teratogenicity. Science 327, 1345–1350 (2010).

7. Chamberlain PP, et al. Structure of the human Cereblon-DDB1-lenalidomide complex reveals basis for responsiveness to thalidomide analogs. Nat Struct Mol Biol 21, 803–809 (2014).

8. Zhu YX, et al. Cereblon expression is required for the antimyeloma activity of lenalidomide and pomalidomide. Blood 118, 4771–4779 (2011).

9. Lopez-Girona A, et al. Cereblon is a direct protein target for immunomodulatory and antiproliferative activities of lenalidomide and pomalidomide. Leukemia 26, 2326–2335 (2012).

10. Lu G, et al. The myeloma drug lenalidomide promotes the cereblon-dependent destruction of Ikaros proteins. Science 343, 305–309 (2014).

11. Kronke J, et al. Lenalidomide causes selective degradation of IKZF1 and IKZF3 in multiple myeloma cells. Science 343, 301–305 (2014).

12. Shaffer AL, et al. IRF4 addiction in multiple myeloma. Nature 454, 226–231 (2008).

13. Matyskiela ME, et al. A Cereblon Modulator (CC-220) with Improved Degradation of Ikaros and Aiolos. J Med Chem 61, 535–542 (2018).

14. Bjorklund CC, et al. Iberdomide (CC-220) is a potent cereblon E3 ligase modulator with antitumor and immunostimulatory activities in lenalidomide- and pomalidomide-resistant multiple myeloma cells with dysregulated CRBN. Leukemia 34, 1197–1201 (2020).

15. Moens CB, Selleri L. Hox cofactors in vertebrate development. Dev Biol 291, 193–206 (2006).

16. Capdevila J, Tsukui T, Rodriquez Esteban C, Zappavigna V, Izpisua Belmonte JC. Control of vertebrate limb outgrowth by the proximal factor Meis2 and distal antagonism of BMPs by Gremlin. Mol Cell 4, 839–849 (1999).

17. Crowley MA, Conlin LK, Zackai EH, Deardorff MA, Thiel BD, Spinner NB. Further evidence for the possible role of MEIS2 in the development of cleft palate and cardiac septum. Am J Med Genet A **152A**, 1326–1327 (2010).

18. Paige SL, et al. A temporal chromatin signature in human embryonic stem cells identifies regulators of cardiac development. Cell 151, 221–232 (2012).

19. Oulad-Abdelghani M, Chazaud C, Bouillet P, Sapin V, Chambon P, Dolle P. Meis2, a novel mouse Pbx-related homeobox gene induced by retinoic acid during differentiation of P19 embryonal carcinoma cells. Dev Dyn 210, 173–183 (1997).

20. Vegi NM, et al. MEIS2 Is an Oncogenic Partner in AML1-ETO-Positive AML. Cell Reports 16, 498–507 (2016).

21. Fischer ES, et al. Structure of the DDB1-CRBN E3 ubiquitin ligase in complex with thalidomide. Nature 512, 49–53 (2014).

22. Ely S, et al. Mutually exclusive cyclin-dependent kinase 4/cyclin D1 and cyclin-dependent kinase 6/cyclin D2 pairing inactivates retinoblastoma protein and promotes cell cycle dysregulation in multiple myeloma. Cancer Res 65, 11345–11353 (2005).

23. Fry DW, et al. Specific inhibition of cyclin-dependent kinase 4/6 by PD 0332991 and associated antitumor activity in human tumor xenografts. Mol Cancer Ther 3, 1427–1438 (2004).

24. Baughn LB, et al. A novel orally active small molecule potently induces G1 arrest in primary myeloma cells and prevents tumor growth by specific inhibition of cyclin-dependent kinase 4/6. Cancer Res 66, 7661–7667 (2006).

25. Huang X, et al. Prolonged early G1 arrest by selective CDK4/CDK6 inhibition sensitizes myeloma cells to cytotoxic killing through cell cycle-coupled loss of IRF4. Blood 120, 1095–1106 (2012).

26. Menu E, et al. A novel therapeutic combination using PD 0332991 and bortezomib: study in the 5T33MM myeloma model. Cancer Res 68, 5519–5523 (2008).

27. Steinebach C, et al. Systematic exploration of different E3 ubiquitin ligases: an approach towards potent and selective CDK6 degraders. Chem Sci 11, 3474–3486 (2020).

28. Ng YLD, et al. Proteomic profiling reveals CDK6 upregulation as a targetable resistance mechanism for lenalidomide in multiple myeloma. Nature Communications 13, 1009 (2022).

29. Tomczak K, Czerwinska P, Wiznerowicz M. The Cancer Genome Atlas (TCGA): an immeasurable source of knowledge. Contemp Oncol (Pozn) 19, A68–77 (2015).

30. T T. A Package for Survival Analysis in R. R package version 3.8-3.) (2024).

31. Settino M, Cannataro M. MMRFBiolinks: an R-package for integrating and analyzing MMRF-CoMMpass data. Brief Bioinform 22, (2021).

32. Martin M. Cutadapt removes adapter sequences from high-throughput sequencing reads. EMBnetjournal 17, 10 (2011).

33. Dobin A, et al. STAR: ultrafast universal RNA-seq aligner. Bioinformatics 29, 15–21 (2013).

34. Trapnell C, et al. Transcript assembly and quantification by RNA-Seq reveals unannotated transcripts and isoform switching during cell differentiation. Nat Biotechnol 28, 511–515 (2010).

35. Roberts A, Trapnell C, Donaghey J, Rinn JL, Pachter L. Improving RNA-Seq expression estimates by correcting for fragment bias. Genome Biol 12, R22 (2011).

36. Subramanian A, et al. Gene set enrichment analysis: a knowledge-based approach for interpreting genome-wide expression profiles. Proc Natl Acad Sci U S A 102, 15545–15550 (2005).

37. Heintel D, et al. High expression of cereblon (CRBN) is associated with improved clinical response in patients with multiple myeloma treated with lenalidomide and dexamethasone. Br J Haematol 161, 695–700 (2013).

38. Roecklein BA, Torok-Storb B. Functionally distinct human marrow stromal cell lines immortalized by transduction with the human papilloma virus E6/E7 genes. Blood 85, 997–1005 (1995).

39. Yan Z, et al. Cdc6 is regulated by E2F and is essential for DNA replication in mammalian cells. Proc Natl Acad Sci U S A 95, 3603–3608 (1998).

40. Kent LN, Leone G. The broken cycle: E2F dysfunction in cancer. Nat Rev Cancer 19, 326–338 (2019).

41. Forbes S, et al. Cosmic 2005. Br J Cancer 94, 318–322 (2006).

42. Escoubet-Lozach L, et al. Pomalidomide and lenalidomide induce p21 WAF-1 expression in both lymphoma and multiple myeloma through a LSD1-mediated epigenetic mechanism. Cancer Res 69, 7347–7356 (2009).

43. Carrano AC, Eytan E, Hershko A, Pagano M. SKP2 is required for ubiquitin-mediated degradation of the CDK inhibitor p27. Nat Cell Biol 1, 193–199 (1999).

44. Lopez-Girona A, et al. Lenalidomide downregulates the cell survival factor, interferon regulatory factor-4, providing a potential mechanistic link for predicting response. Br J Haematol 154, 325–336 (2011).

45. Fedele PL, et al. IMiDs prime myeloma cells for daratumumab-mediated cytotoxicity through loss of Ikaros and Aiolos. Blood 132, 2166–2178 (2018).

46. Chen Q, et al. Apo2L/TRAIL and Bcl-2-related proteins regulate type I interferon-induced apoptosis in multiple myeloma. Blood 98, 2183–2192 (2001).

47. Yang Y, et al. Exploiting synthetic lethality for the therapy of ABC diffuse large B cell lymphoma. Cancer Cell 21, 723–737 (2012).

48. Ursini-Siegel J, et al. TRAIL/Apo-2 ligand induces primary plasma cell apoptosis. J Immunol 169, 5505–5513 (2002).

49. Hatzoglou A, et al. TNF receptor family member BCMA (B cell maturation) associates with TNF receptor-associated factor (TRAF) 1, TRAF2, and TRAF3 and activates NF-kappa B, elk-1, c-Jun N-terminal kinase, and p38 mitogen-activated protein kinase. J Immunol 165, 1322–1330 (2000).

50. Novak AJ, et al. Expression of BCMA, TACI, and BAFF-R in multiple myeloma: a mechanism for growth and survival. Blood 103, 689–694 (2004).

51. Abruzzese MP, et al. The homeobox transcription factor MEIS2 is a regulator of cancer cell survival and IMiDs activity in Multiple Myeloma: modulation by Bromodomain and Extra-Terminal (BET) protein inhibitors. Cell Death & Disease 10, 324 (2019).

52. Agoston Z, Schulte D. Meis2 competes with the Groucho co-repressor Tle4 for binding to Otx2 and specifies tectal fate without induction of a secondary midbrain-hindbrain boundary organizer. Development 136, 3311–3322 (2009).

53. Conte I, et al. miR-204 is required for lens and retinal development via Meis2 targeting. Proc Natl Acad Sci U S A 107, 15491–15496 (2010).

54. Heine P, Dohle E, Bumsted-O’Brien K, Engelkamp D, Schulte D. Evidence for an evolutionary conserved role of homothorax/Meis1/2 during vertebrate retina development. Development 135, 805–811 (2008).

55. Yakushiji-Kaminatsui N, et al. RING1 proteins contribute to early proximal-distal specification of the forelimb bud by restricting Meis2 expression. Development 143, 276–285 (2016).

56. Agoston Z, et al. Meis2 is a Pax6 co-factor in neurogenesis and dopaminergic periglomerular fate specification in the adult olfactory bulb. Development 141, 28–38 (2014).

57. Bjerke GA, Hyman-Walsh C, Wotton D. Cooperative transcriptional activation by Klf4, Meis2, and Pbx1. Mol Cell Biol 31, 3723–3733 (2011).

58. Bumsted-O’Brien KM, Hendrickson A, Haverkamp S, Ashery-Padan R, Schulte D. Expression of the homeodomain transcription factor Meis2 in the embryonic and postnatal retina. J Comp Neurol 505, 58–72 (2007).

59. Crijns AP, et al. MEIS and PBX homeobox proteins in ovarian cancer. Eur J Cancer 43, 2495–2505 (2007).

60. Zha Y, et al. MEIS2 is essential for neuroblastoma cell survival and proliferation by transcriptional control of M-phase progression. Cell Death & Disease 5, e1417–e1417 (2014).

61. Zhou P, Howley PM. Ubiquitination and degradation of the substrate recognition subunits of SCF ubiquitin-protein ligases. Mol Cell 2, 571–580 (1998).

62. Goel S, et al. CDK4/6 inhibition triggers anti-tumour immunity. Nature 548, 471–475 (2017).

63. Deng J, et al. CDK4/6 Inhibition Augments Antitumor Immunity by Enhancing T-cell Activation. Cancer Discov 8, 216–233 (2018).

64. Jerby-Arnon L, et al. A Cancer Cell Program Promotes T Cell Exclusion and Resistance to Checkpoint Blockade. Cell 175, 984–997 e924 (2018).

65. Lelliott EJ, et al. CDK4/6 inhibition promotes anti-tumor immunity through the induction of T cell memory. Cancer Discovery, candisc.1554.2020 (2021).

66. Heckler M, et al. Inhibition of CDK4/6 promotes CD8 T-cell memory formation. Cancer Discov, (2021).

67. Tam CS, et al. Ibrutinib plus Venetoclax for the Treatment of Mantle-Cell Lymphoma. N Engl J Med 378, 1211–1223 (2018).

68. Niesvizky R, et al. Phase 1/2 study of cyclin-dependent kinase (CDK)4/6 inhibitor palbociclib (PD-0332991) with bortezomib and dexamethasone in relapsed/refractory multiple myeloma. Leuk Lymphoma 56, 3320–3328 (2015).

