## Supplementary material for "CDK4/6 Inhibition Reverses MEIS2 Suppression of CRL4^CRBN^ to Enhance Immunomodulatory Drug Therapy in Multiple Myeloma": 251014 MEIS2 mspt supplement.pdf

Running title: CDK4/6i reverses MEIS2 inhibition of CRL4$^{CBRN}$

Xiangao Huang[1,2], David Jayabalan[1,3], Maurizio Di Liberto[1,2], Zhengming Chen[4], Scott A. Ely[1], Tomer M. Mark[3], Gang Lin[2,5], Ruben Niesvizky[2,3,*] and Selina Chen-Kiang[1,2,6,*]

[1]Department of Pathology and Laboratory Medicine, [2]Sandra and Edward Meyer Cancer Center [3]Department of Medicine, [4]Department of Public Health, [5]Department of Microbiology and Immunology, and [6]Graduate Program in Immunology and Microbial Pathogenesis, Weill Cornell Medicine, New York, NY 10065.

*Present address for Scott Ely:* Memorial Sloan *Kettering Cancer Center,* New York, NY 10065

*Present address for Tomer M. Mark*: Karyopharm Therapeutics, Newton, MA 02459

*Co-*corresponding authors:*

Selina Chen-Kiang

Department of Pathology and Laboratory Medicine, Weill Cornell Medicine, 1300 York Avenue New York, NY, 10065. Tel: 212-746-6440.

Ruben Niesvizky

Department of Medicine, Weill Cornell Medicine, 425 East 61$^{st}$ Street, New York, New York 10065. Tel: 646-962-6576.

**Supplementary information includes:**

Supplementary Figures 1-5

Supplementary Tables 1, 2 and 4

**Supplementary Figures**

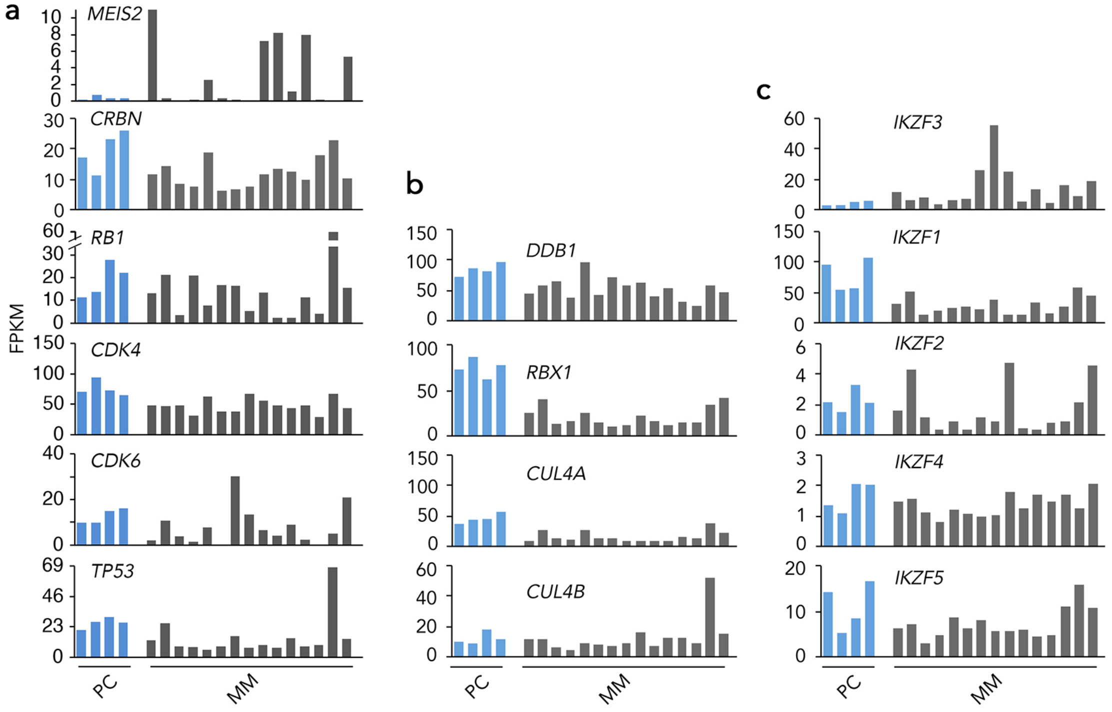

Supplementary Fig. 1

**Supplementary Fig. 1. Expression of cell cycle genes.** (A-C) RNA-Seq analysis of mRNA abundance (FPKM) of indicated genes in normal bone marrow plasma cells (PC), and freshly isolated primary bone marrow myeloma cells (MM).

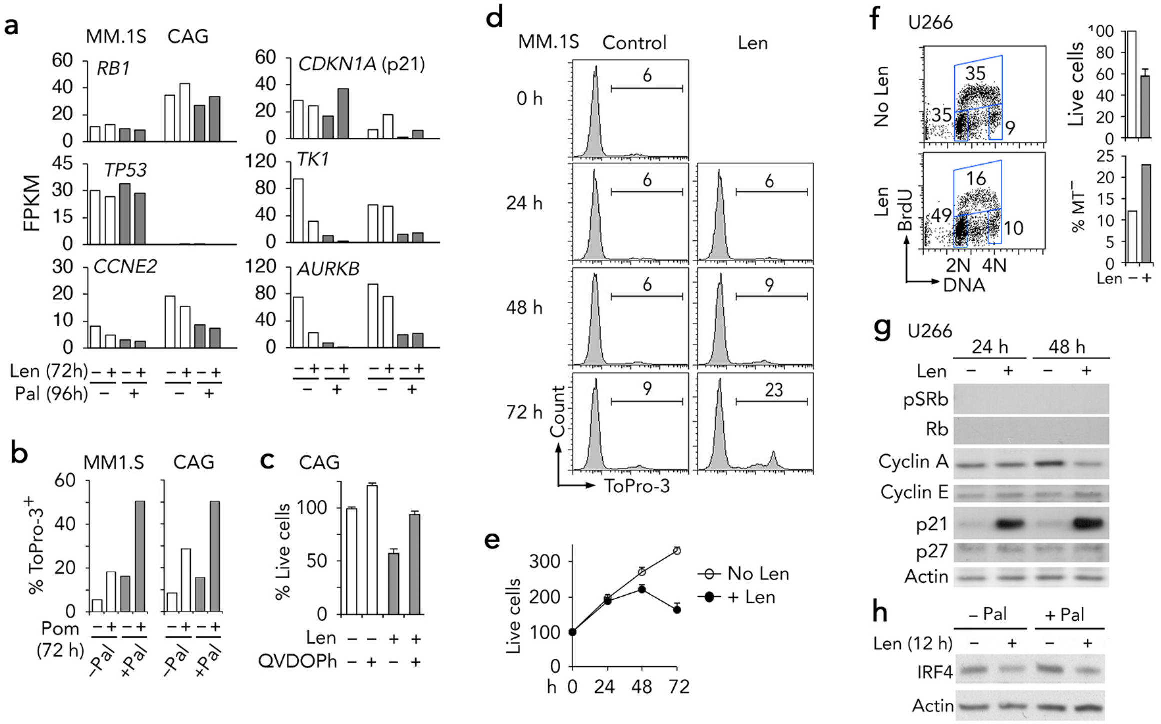

Supplementary Fig. 2

**Supplementary Fig. 2. Lenalidomide induced cell death and inhibited cell proliferation independent of Rb protein.** (A) RNA-seq analysis of mRNA abundance of indicated genes in MM.1S and CAG cells treated with Pal and Len for time indicated. (B) Cell death in MM.1S and CAG cells cultured with pomalidomide (Pom, 3μM) for 72 hours +/- Pal pretreatment. (C) Percentage of live CAG cells exposed to Len for 72 hours in the presence or absence of the caspase inhibitor Q-VD-OPh (20 μM). (D) Analysis of cell death (ToPro-3 staining) in MM.1S cells cultured with lenalidomide (Len, 3 μM ) for time indicated. (E) Analysis of total live cells (trypan blue exclusion assay) in MM.1S cells treated with Len for time indicated. (F) Analysis of cell cycle, total live cells, and cell death (MitoTracker Red staining) in U266 cells cultured with Len for 72 hours. (G) Immunoblotting of indicated proteins in U266 cells treated with Len for time indicated. (H) Immunoblotting of IRF4 in U266 exposed to Len for 12 hours +/- Pal pretreatment.

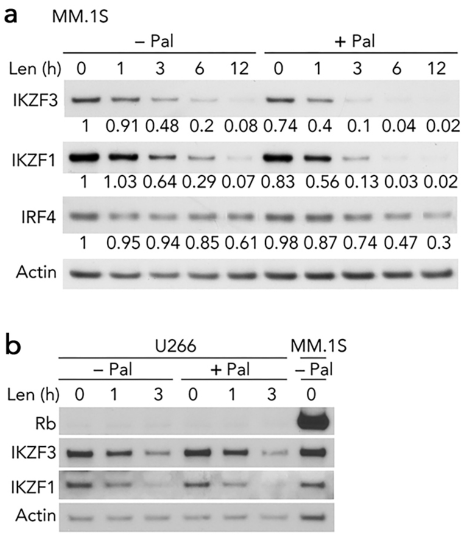

Supplementary Fig. 3

**Supplementary Fig. 3. Combination of Pal and Len in myeloma cell lines.** (A-B) Immunoblotting analysis of indicated proteins in MM.1S and U266 cells cultured with Len for time indicated +/- Pal pretreatment. Levels of IKZF3, IKZF1 and IRF4 proteins were normalized to Actin and calculated against the untreated control (lane 1).

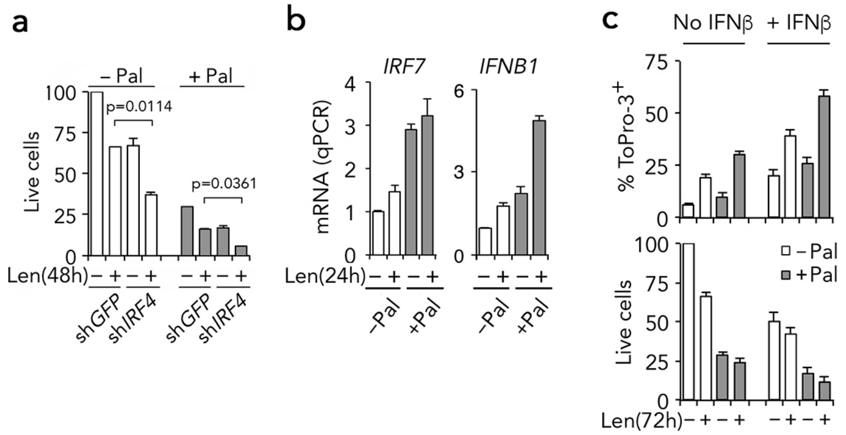

Supplementary Fig. 4

**Supplementary Fig. 4. Gene expression of interferon-regulatory factors in myeloma cells.** (A) Viability of KMS12PE cells infected with GFP shRNA (shGFP) or IRF4-shRNA (shIRF4) lentivirus for 24 hours and cultured with Len for 48 hours +/- Pal pretreatment. (B) q-PCR analysis of IRF7 and IFNB1 in MM.1S cells exposed to Len for 24 hours +/- Pal pretreatment. (C) Percentages of ToPro-3+ MM.1S cells and total live cells treated with Len for 72 hours +/- Pal pretreatment in the presence or absence of IFNβ (500 ng/ml).

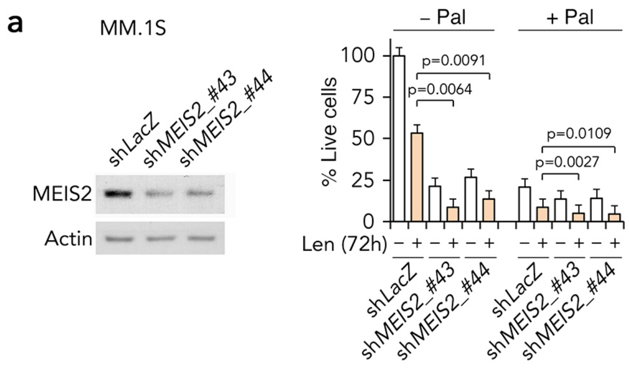

**b**

Top 10 gene sets enriched by MEIS2 knockdown

| Gene Set | SIZE | NES | NOM p-val |
|---|---|---|---|
| SANA_RESPONSE_TO_IFNG_UP | 37 | 2.191 | <0.001 |
| NIKOLSKY_BREAST_CANCER_16P13_AMPLIC | 56 | 2.109 | <0.001 |
| REACTOME_INTERFERON_ALPHA_BETA_SIG | 26 | 2.104 | <0.001 |
| REACTOME_GLYCOSPHINGOLIPID_METABO | 18 | 2.042 | 0.0023 |
| REACTOME_INTERFERON_GAMMA_SIGNAL | 31 | 2.015 | <0.001 |
| BROCKE_APOPTOSIS_REVERSED_BY_IL6 | 101 | 2.011 | <0.001 |
| MANALO_HYPOXIA_UP | 65 | 1.984 | <0.001 |
| ONGUSAHA_TP53_TARGETS | 20 | 1.969 | 0.0025 |
| SUBTIL_PROGESTIN_TARGETS | 20 | 1.958 | <0.001 |
| BROWNE_INTERFERON_RESPONSIVE_GENE | 32 | 1.953 | 0.0024 |

Supplementary Fig. 5

**Supplementary Fig. 5. Gene sets enriched by MEIS2 knockdown**. (A) Immunoblotting of MEIS2 in MM.1S infected with *LacZ-29* shRNA *(*sh*LacZ)*, or sh*MEIS2*_#43 (TRCN0000016043) or sh*MEIS*2_#44 (TRCN0000016044) lentivirus. Cell viability was determined after 72 h of Len exposure +/- Pal pretreatment (bottom). (B) GSEA identified top 10 gene sets enriched by MEIS2 knockdown in MM.1S cells.

### Supplementary Tables

**Supplemental Table 1. Multiple Myeloma Cases in This Study**

| MM # | Age/Sex | Ig isotype | Stage | % Ki67⁺/CD138⁺ | # Prior therapy | Post-biopsy IMiD therapy | Best response | IMiD response *ex vivo* | Figure No. |
|---|---|---|---|---|---|---|---|---|---|
| 1 | 78/F | Gκ | III | 13.5 | 5 | ClaPd | VGPR | Yes | 2D-E,2F |
| 18 | 62/F | Aλ | II | 4 | 0 | RVD | CR | Yes | 2F,5A-D |
| 19 | 56/F | Aλ | III | 23 | 3 | RD | VGPR | Yes | 2F,5A-D |
| 10 | 57/M | Gκ | II | 14 | 0 | CBiRD | VGPR | Yes | 2F,3A, 5A-D |
| 11 | 64/M | Gκ | II | NA | 0 | CBiRD | VGPR | Yes | 2F,5A-D |
| 2 | 56/F | Gκ | III | 8 | 9 | ClaPd | SD | Yes | 2F |
| 3 | 55/F | Gκ | III | 1 | 7 | ClaPd | PR | Yes | 2F |
| 5 | 41/F | Aλ | III | 10 | 8 | ClaPd | VGPR | Yes | 2F |
| 6 | 77/F | Aλ | II | 1 | 0 | NA | NA | Yes | 2F |
| 8 | 67/M | Aλ | II | 1 | 0 | NA | NA | Yes | 2F |
| 9 | 79/M | Gκ | III | 30 | 5 | ClaPd | SD | Yes | 2F |
| 21 | 63/M | Gκ | II | NA | 3 | R(HD)-ASCT | CR | Yes | 2F |
| 31 | 52/F | Gκ | II | 4 | 0 | CBiRD | PR | Yes | 2F,4F |
| 12 | 64/F | Gκ | II | NA | 0 | RD | PR | Yes | 2F |
| 16 | 61/M | Gκ | II | 11 | 1 | RD | CR | Yes | 2H |
| 17 | 46/M | Aλ | III | 1 | 6 | ClaPd | SD | Yes | 2H |
| 22 | 67/M | Gκ | I | NA | 0 | NA | NA | NA | 4E |
| 24 | 54/F | Gλ | II | NA | 0 | NA | NA | NA | 1C |
| 25 | 82/F | Gκ | II | NA | 0 | NA | NA | NA | 1C |
| 27 | 82/M | Gκ | III | NA | 2 | NA | NA | NA | 1C |
| 28 | 67/M | κ | III | NA | 6 | ClaPd | SD | NA | 1C |
| 29 | 71/F | λ | II | NA | 0 | CBiRD | VGPR | NA | 4G |

| MM # | Age/Sex | Ig isotype | Stage | % Ki67⁺/CD138⁺ | # Prior therapy | Before-biopsy IMiD therapy | Best response | IMiD response *ex vivo* | Figure No. |
|---|---|---|---|---|---|---|---|---|---|
| 15 | 73/F | Gκ | II | NA | 3 | RVD | PD | No | 2G-H |
| 20 | 66/F | Aκ | II | 0 | 3 | R | PD | No | 2G,5A-D |
| 30 | 44/M | Aλ | III | 1 | 3 | RVD | PD | No | 2G,5A-D |
| 13 | 71/M | Aκ | II | 5 | 4 | RD | PD | No | 2G |
| 14 | 74/F | Gκ | II | 24 | 4 | R | Toxicity | No | 2G |

| MM # | Age/Sex | Ig isotype | Stage | % Ki67⁺/CD138⁺ | # Prior therapy | Post-biopsy IMiD therapy | Best response | IMiD response *ex vivo* | Figure no. |
|---|---|---|---|---|---|---|---|---|---|
| 32 | 57/F | Gκ | I | 0.5 | 0 | RVD | PR | No | 2B-C |
| 33 | 75/M | Gκ | I | 1 | 0 | NA | NA | No | 2C |
| 34 | 60/F | κ | III | 7 | 12 | NA | NA | No | 2C |

**Abbreviations**: Bi, Biaxin; C, carfilzomib; Cla, clarithromycin; D, dexamethasone; Ig, immunoglobulin; P, pomalidomide; R, Revlimid (lenalidomide); V, Velcade (bortezomib); NA, data not available; Pt, patient.

**Stage**: Durie-Salmon staging

**Therapy**: R - Revlimid; RD - Revlimid and dexamethasone; ClaPd - clarithromycin, pomalidomide and dexamethasone; R(HD)-ASCT - high dose Revlimid-autologous stem cell transplant; CBiRD - carfilzomib, Biaxin, Revlimid and dexamethasone; RVD - Revlimid, Velcade and dexamethasone.

**Response**: CR, complete response (100% reduction of M-Spike, the monoclonal Ig characteristic of myeloma); VGPR, very good partial response (≥ 90% reduction of M-Spike); PR, partial response (≥50% reduction of M-Spike); SD, stable disease (≤ 25 % reduction of M-Spike); PD, progression disease.

**Supplementary Table 2.** The pairwise-comparison p-values determined by two-way ANOVA analysis of viable primary myeloma cells treated with Pal in combination with lenalidomide or pomalidomide

| MM | 1 | 18 | 19 | 10 | 11 | 2 | 3 |
|---|---|---|---|---|---|---|---|
| Len vs Pal+Len | 0.0096 | 0.043 | 0.017 | 0.0285 | 0.0473 | 0.0755 | 0.0199 |
| Pal vs Pal+Len | 0.0009 | <0.0001 | 0.0012 | 0.0013 | 0.0001 | 0.0015 | 0.0046 |
| MM | 5 | 6 | 8 | 9 | 21 | 31 | 12 |
| Len vs Pal+Len | 0.009 | 0.1588 | <0.0001 | 0.0005 | 0.0042 | 0,0001 | 0.9654 |
| Pal vs Pal+Len | 0.0923 | 0.0195 | <0.0001 | 0.0002 | 0.0004 | 0.0001 | <0.0001 |
| MM | 15 | 20 | 30 | 13 | 14 | | |
| Len vs Pal+Len | 0.6161 | 0.947 | 0.0013 | 0.6359 | 0.2638 | | |
| Pal vs Pal+Len | 0.6186 | 0.9728 | 0.7525 | 0.5701 | 0.9246 | | |
| MM | 9 | 16 | 17 | 15 | | | |
| Pom vs Pal+Len | 0.0053 | <0.0001 | 0.0065 | 0.8724 | | | |
| Pal vs Pal+Len | <0.0001 | <0.0001 | 0.0001 | 0.602 | | | |

Abbreviations: MM, primary BM myeloma cells; Pal, palbociclib; Len, lenalidomide; Pom, pomalidomide.

**Supplementary Table 4.** Gene sets enriched in BMMCs in response to lenalidomide (Len), palbociclib (Pal) or Len plus Pal (PaLen).

| Dataset | Gene Set | SIZE | NES | NOM p-val | FDR q-val |
|---|---|---|---|---|---|
| Group I/Group II in response to Len | NUYTTEN_NIPP1_TARGETS_UP | 27 | 2.00 | <0.0001 | 0.0176 |
| | BROWNE_INTERFERON_RESPONSIVE_GENES | 19 | 1.98 | <0.0001 | 0.0123 |
| | TAKEDA_TARGETS_OF_NUP98_HOXA9_FUSION_3D_UP | 29 | 1.91 | <0.0001 | 0.0137 |
| | TAKEDA_TARGETS_OF_NUP98_HOXA9_FUSION_10D_UP | 28 | 1.90 | <0.0001 | 0.0116 |
| | FULCHER_INFLAMMATORY_RESPONSE_LECTIN_VS_LPS_DN | 30 | 1.86 | <0.0001 | 0.0147 |
| | TAKEDA_TARGETS_OF_NUP98_HOXA9_FUSION_16D_UP | 25 | 1.79 | 0.0013 | 0.0200 |
| | HELLER_SILENCED_BY_METHYLATION_UP | 22 | 1.83 | 0.0013 | 0.0159 |
| | RADAEVA_RESPONSE_TO_IFNA1_UP | 15 | 1.83 | 0.0014 | 0.0175 |
| | MOSERLE_IFNA_RESPONSE | 18 | 1.94 | 0.0014 | 0.0131 |
| | TAKEDA_TARGETS_OF_NUP98_HOXA9_FUSION_8D_UP | 21 | 1.75 | 0.0027 | 0.0265 |
| | NUYTTEN_EZH2_TARGETS_UP | 42 | 1.77 | 0.0035 | 0.0254 |
| | HECKER_IFNB1_TARGETS | 23 | 1.74 | 0.0039 | 0.0254 |
| | DEBIASI_APOPTOSIS_BY_REOVIRUS_INFECTION_UP | 17 | 1.91 | 0.0039 | 0.0140 |
| | BOSCO_INTERFERON_INDUCED_ANTIVIRAL_MODULE | 21 | 1.87 | 0.0054 | 0.0128 |
| | DAUER_STAT3_TARGETS_DN | 20 | 1.75 | 0.0066 | 0.0249 |
| | SANA_RESPONSE_TO_IFNG_UP | 15 | 1.80 | 0.0080 | 0.0206 |
| | BENPORATH_ES_WITH_H3K27ME3 | 17 | 1.58 | 0.0188 | 0.1004 |
| | GRAESSMANN_RESPONSE_TO_MC_AND_SERUM_DEPRIVATIO | 19 | 1.56 | 0.0221 | 0.1063 |
| | ZHOU_INFLAMMATORY_RESPONSE_LPS_UP | 20 | 1.58 | 0.0246 | 0.0976 |
| | HELLER_HDAC_TARGETS_SILENCED_BY_METHYLATION_UP | 17 | 1.58 | 0.0258 | 0.1058 |
| | DODD_NASOPHARYNGEAL_CARCINOMA_DN | 20 | 1.57 | 0.0305 | 0.0979 |
| | GRAESSMANN_APOPTOSIS_BY_DOXORUBICIN_UP | 37 | 1.49 | 0.0323 | 0.1507 |
| | REACTOME_INTERFERON_ALPHA_BETA_SIGNALING | 16 | 1.53 | 0.0353 | 0.1220 |
| | GRAESSMANN_APOPTOSIS_BY_SERUM_DEPRIVATION_UP | 28 | 1.53 | 0.0391 | 0.1255 |

| Dataset | Gene Set | SIZE | NES | NOM p-val | FDR q-val |
|---|---|---|---|---|---|
| Group I/Group II in response to Pal | TAKEDA_TARGETS_OF_NUP98_HOXA9_FUSION_8D_UP | 21 | 1.74 | <0.0001 | 0.1024 |
| | TAKEDA_TARGETS_OF_NUP98_HOXA9_FUSION_3D_UP | 29 | 1.67 | 0.0011 | 0.0931 |
| | TAKEDA_TARGETS_OF_NUP98_HOXA9_FUSION_10D_UP | 28 | 1.61 | 0.0021 | 0.0646 |
| | FULCHER_INFLAMMATORY_RESPONSE_LECTIN_VS_LPS_DN | 30 | 1.63 | 0.0031 | 0.0750 |
| | NUYTTEN_NIPP1_TARGETS_UP | 27 | 1.60 | 0.0042 | 0.0674 |
| | BROWNE_INTERFERON_RESPONSIVE_GENES | 19 | 1.61 | 0.0043 | 0.0708 |
| | HELLER_HDAC_TARGETS_SILENCED_BY_METHYLATION_UP | 17 | 1.67 | 0.0044 | 0.1396 |
| | HECKER_IFNB1_TARGETS | 23 | 1.63 | 0.0054 | 0.0702 |
| | DAUER_STAT3_TARGETS_DN | 20 | 1.64 | 0.0064 | 0.0845 |
| | BOSCO_INTERFERON_INDUCED_ANTIVIRAL_MODULE | 21 | 1.64 | 0.0076 | 0.0975 |
| | RADAEVA_RESPONSE_TO_IFNA1_UP | 15 | 1.59 | 0.0124 | 0.0701 |
| | HELLER_SILENCED_BY_METHYLATION_UP | 22 | 1.54 | 0.0193 | 0.1114 |
| | MOSERLE_IFNA_RESPONSE | 18 | 1.49 | 0.0195 | 0.1513 |
| | SENESE_HDAC3_TARGETS_UP | 18 | 1.49 | 0.0240 | 0.1617 |
| | DEBIASI_APOPTOSIS_BY_REOVIRUS_INFECTION_UP | 17 | 1.52 | 0.0268 | 0.1316 |
| | TAKEDA_TARGETS_OF_NUP98_HOXA9_FUSION_16D_UP | 25 | 1.44 | 0.0298 | 0.1833 |
| | REACTOME_INTERFERON_SIGNALING | 22 | 1.46 | 0.0302 | 0.1888 |
| | WALLACE_PROSTATE_CANCER_RACE_UP | 33 | 1.41 | 0.0352 | 0.2220 |
| | BENPORATH_EED_TARGETS | 16 | 1.46 | 0.0356 | 0.1780 |
| | REACTOME_CYTOKINE_SIGNALING_IN_IMMUNE_SYSTEM | 23 | 1.45 | 0.0362 | 0.1863 |
| | REACTOME_INTERFERON_ALPHA_BETA_SIGNALING | 16 | 1.46 | 0.0435 | 0.1769 |
| | GOZGIT_ESR1_TARGETS_DN | 28 | 1.43 | 0.0482 | 0.1896 |

| Dataset | Gene Set | SIZE | NES | NOM p-val | FDR q-val |
|---|---|---|---|---|---|

| | | | | | |
|---|---|---|---|---|---|
| Group I/Group II in response to PaLen | NUYTTEN_NIPP1_TARGETS_UP | 27 | 2.33 | <0.0001 | 0.0000 |
| | TAKEDA_TARGETS_OF_NUP98_HOXA9_FUSION_3D_UP | 29 | 2.18 | <0.0001 | 0.0000 |
| | BROWNE_INTERFERON_RESPONSIVE_GENES | 19 | 2.15 | <0.0001 | 0.0007 |
| | DEBIASI_APOPTOSIS_BY_REOVIRUS_INFECTION_UP | 17 | 2.14 | <0.0001 | 0.0005 |
| | FULCHER_INFLAMMATORY_RESPONSE_LECTIN_VS_LPS_DN | 30 | 2.13 | <0.0001 | 0.0004 |
| | NUYTTEN_EZH2_TARGETS_UP | 42 | 2.12 | <0.0001 | 0.0005 |
| | HECKER_IFNB1_TARGETS | 23 | 2.10 | <0.0001 | 0.0006 |
| | RADAEVA_RESPONSE_TO_IFNA1_UP | 15 | 2.04 | <0.0001 | 0.0015 |
| | BOSCO_INTERFERON_INDUCED_ANTIVIRAL_MODULE | 21 | 2.03 | <0.0001 | 0.0014 |
| | TAKEDA_TARGETS_OF_NUP98_HOXA9_FUSION_10D_UP | 28 | 2.01 | <0.0001 | 0.0015 |
| | DAUER_STAT3_TARGETS_DN | 20 | 2.01 | <0.0001 | 0.0015 |
| | MOSERLE_IFNA_RESPONSE | 18 | 1.98 | <0.0001 | 0.0019 |
| | TAKEDA_TARGETS_OF_NUP98_HOXA9_FUSION_8D_UP | 21 | 1.91 | <0.0001 | 0.0051 |
| | SANA_RESPONSE_TO_IFNG_UP | 15 | 1.91 | 0.0017 | 0.0053 |
| | HELLER_HDAC_TARGETS_SILENCED_BY_METHYLATION_UP | 17 | 1.86 | 0.0033 | 0.0083 |
| | REACTOME_INTERFERON_ALPHA_BETA_SIGNALING | 16 | 1.94 | 0.0033 | 0.0034 |
| | REACTOME_INTERFERON_SIGNALING | 22 | 1.80 | 0.0049 | 0.0141 |
| | HELLER_SILENCED_BY_METHYLATION_UP | 22 | 1.86 | 0.0051 | 0.0084 |
| | GOZGIT_ESR1_TARGETS_DN | 28 | 1.65 | 0.0067 | 0.0450 |
| | DODD_NASOPHARYNGEAL_CARCINOMA_DN | 20 | 1.69 | 0.0081 | 0.0340 |
| | TAKEDA_TARGETS_OF_NUP98_HOXA9_FUSION_16D_UP | 25 | 1.74 | 0.0098 | 0.0254 |
| | GRAESSMANN_APOPTOSIS_BY_DOXORUBICIN_UP | 37 | 1.65 | 0.0110 | 0.0447 |
| | ZHOU_INFLAMMATORY_RESPONSE_LPS_UP | 20 | 1.72 | 0.0128 | 0.0277 |
| | MARKEY_RB1_ACUTE_LOF_DN | 16 | 1.64 | 0.0209 | 0.0462 |
| | REACTOME_CYTOKINE_SIGNALING_IN_IMMUNE_SYSTEM | 23 | 1.61 | 0.0213 | 0.0541 |
| | GRAESSMANN_RESPONSE_TO_MC_AND_SERUM_DEPRIVATIO | 19 | 1.57 | 0.0231 | 0.0713 |
| | REACTOME_IMMUNE_SYSTEM | 43 | 1.49 | 0.0375 | 0.1038 |
| | GRAESSMANN_APOPTOSIS_BY_SERUM_DEPRIVATION_UP | 28 | 1.50 | 0.0442 | 0.1046 |

Notes: Gene set enrichment analysis (GSEA) identified gene sets that were specifically enriched by Len, Pal or PaLen as determined by the normalized enrichment score (NES) with a nominal (NOM) p-value < 0.05.
